# Quality assessment of single-cell RNA sequencing data by coverage skewness analysis

**DOI:** 10.1101/2019.12.31.890269

**Authors:** Imad Abugessaisa, Shuhei Noguchi, Melissa Cardon, Akira Hasegawa, Kazuhide Watanabe, Masataka Takahashi, Harukazu Suzuki, Shintaro Katayama, Juha Kere, Takeya Kasukawa

## Abstract

Analysis and interpretation of single-cell RNA-sequencing (scRNA-seq) experiments are compromised by the presence of poor quality cells. For meaningful analyses, such poor quality cells should be excluded to avoid biases and large variation. However, no clear guidelines exist. We introduce SkewC, a novel quality-assessment method to identify poor quality single-cells in scRNA-seq experiments. The method is based on the assessment of gene coverage for each single cell and its skewness as a quality measure. To validate the method, we investigated the impact of poor quality cells on downstream analyses and compared biological differences between typical and poor quality cells. Moreover, we measured the ratio of intergenic expression, suggesting genomic contamination, and foreign organism contamination of single-cell samples. SkewC is tested in 37,993 single-cells generated by 15 scRNA-seq protocols. We envision SkewC as an indispensable QC method to be incorporated into scRNA-seq experiment to preclude the possibility of scRNA-seq data misinterpretation.

---

Recent advances in scRNA-seq methods and protocols have enabled new discoveries and insights in the biology of cells^[1, 2]^. The method has been used to profile gene expression of individual cells under different biological conditions, and to identify new cell types and provide knowledge about different biological processes^[3]^.

Data quality measures and quality-control (QC) methods aim to provide confidence in the quality of the dataset and assure the robustness, reproducibility and high quality of any experimental study. In bulk RNA-seq experiments, different data quality measures are applied at consecutive experimental stages. At the early stage of experiment, the integrity of RNA (RIN value) is measured, the quality of raw sequences is evaluated (FASTQ) as well as the quality of the alignments (MAPQ), sequencing depth, and expression level.

In scRNA-seq some standard measures cannot be used, and the data quality may vary highly due to variation for biological reasons or experimental procedures. The variation of experimental procedures includes cell capture methods, target of the sequencing protocols, and reaction failure, to name just a few. Cell capture method might expose individual cells to stress and cause cell death. Cell capture site may contain debris due to broken cells or contain multiple cells instead of a single cell. RNA-seq protocols are designed to capture reads either at the end of the gene (5’or 3’ end) or the full gene body (entire transcript). The multitude of the scRNA-seq methods increases the complexity on the required quality assessment of the resulting dataset. The failure or inadequate quality assessment might lead to the presence of poor quality cells (dead or live cells^[4]^) and thus incorrect interpretation or compromised resolution, resulting from mis-clustering errors, propagation of specific cell type population, or poor sensitivity to detect differentially expressed genes (DEGs). A worst-case scenario in the classification of cells is misinterpretation of a cluster of the poor-quality cells as a new cell type.

As reported in several publications ^[4-9]^, identification of poor quality cells is challenging since they might represent a large population of cells and may not be limited to dead cells. To identify poor quality cells experimentally (e.g., microscopic techniques or cell staining), is laborious and involve further manipulations possibly affecting the transcriptome. Computationally, several tools and methods have been developed to identify poor quality cells [5-7, 9]. The first group of the methods classify the cells based on resulting sequence read counts, number of expressed genes, gene expression patterns to detect outliers or cutoff value based on library size [4, 7, 9]. The second group of computational methods use machine learning techniques (classifier-supervised learning) to classify cells based on their normalized expression profile and a training set. The training dataset is generated from experimentally classified scRNA-seq data ^[6]^. A full list of the current tools for single-cell QC is available at ^[10]^.

The challenge is to develop a QC method specific to scRNA-seq and applicable to different types of scRNA-seq data. The above methods are based on existing approaches used in bulk RNA-seq QC and analysis, and they ignore the characteristics of scRNA-seq experiment, variation of methodology and the quality properties of individual cells compared to a bulk RNA-seq sample, resulting in limitations both in terms of the classification result or implementation^[6]^. Here we considered such limitations and introduced new approaches able to segregate poor quality cells from the good quality cells. SkewC enabled us to identify two classes of single cells that we refer to as typical cells with prototypical coverage profile and skewed cells with a skewed coverage profile.

## Results

### Wide discrepancies in gene body coverage among scRNA-seq protocols

Gene body (full transcript length) coverage considers the distribution of the sequence tags over the entire transcripts. We computed and analyzed the gene body coverage of different protocols (**Online Methods**). Remarkably, the gene body coverage shows wide differences among the data-sets generated by different scRNA-seq protocols (**Fig. 1a and b**; **Panels a and b in Supplementary Figs. 1-6**). This indicates major variation and differences in scRNA-seq datasets produced by different protocols.

**Figure 1:**
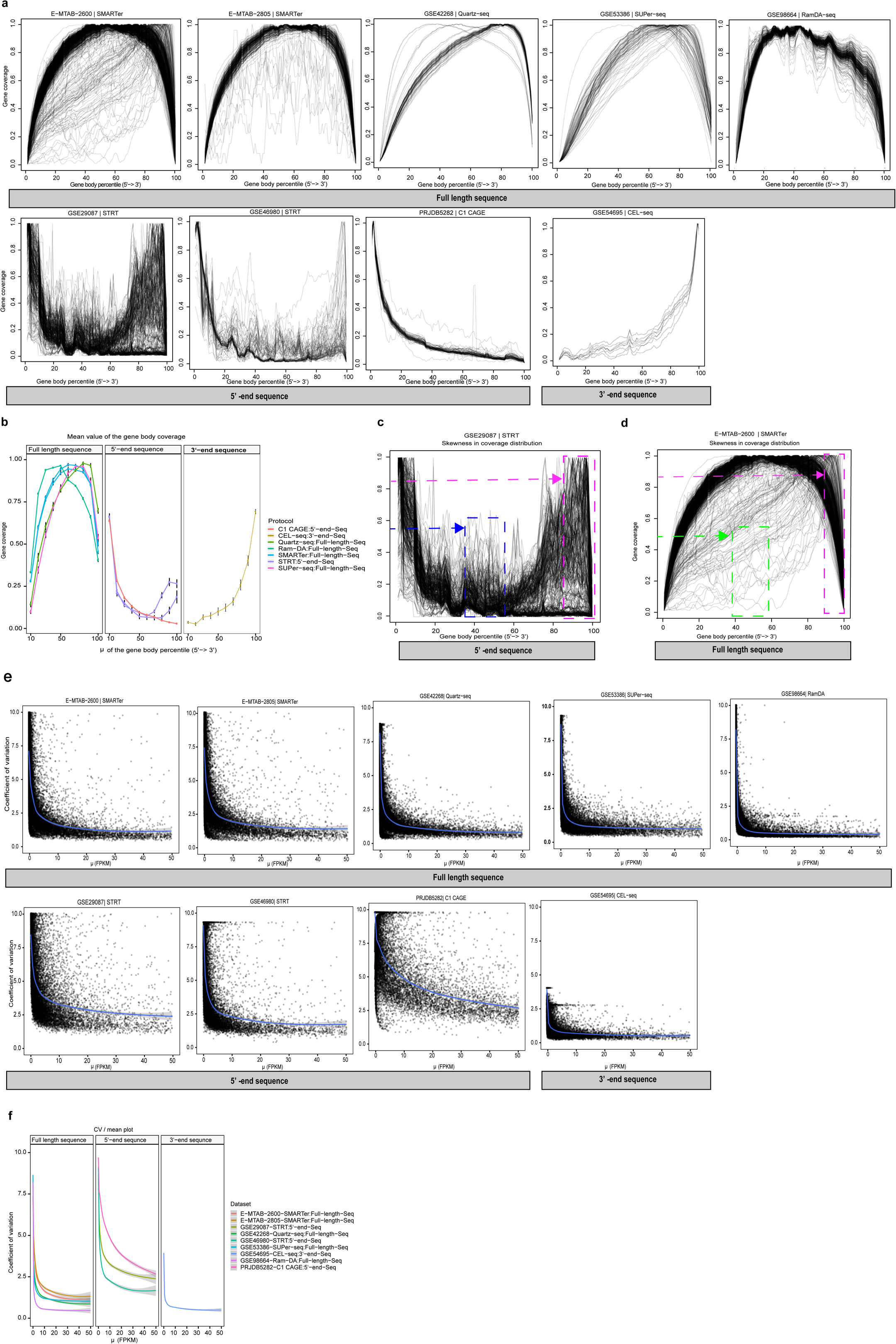
Gene coverage skewness and variation in expression among single-cell RNA-Seq protocols using mESCs. 9 datasets from mouse embryonic stem cells. (a) Distribution of the mapped reads (tags) across the genes. Each panel shows gene body coverage percentile per dataset. The x-axis represents the gene body from 5’ end to 3’ end scaled from 0-100, and the y-axis gene coverage (0-1). Each line represents a single cell. (b) Mean calculated for bin size = 10. (c-d) Skewed distribution of the gene body coverage. (c) & (d) show the bias towards the 3’-end of the gene body (magenta dashed box) and low coverage in the middle of the gene body (blue dashed box). (STRT as 5’-end sequence protocol) shows the bias towards the 3’-end of the gene body (magenta dashed box) and high coverage in the middle of the gene body (green dashed box). (e) The variability in gene expression plot. The X-axis is the mean of the normalized gene expression (FPKM), Y-Axis is the coefficient of variation. CV/mean correlates the sequence depths and variability in gene expression among different protocols. Figure (f), grouping of the smooth lines from (e).

We investigated the pattern of the gene body coverage for each single cell in individual datasets. The visualization of the gene body coverage profile revealed two patterns of gene body coverage. The first set of single-cells show well clustered gene coverage distribution according to the target sequence of the protocol. The second set of single-cells showed skewed gene body coverage distributions. The skewness in the distribution could be observed in term of coverage bias towards specific gene region.

In one type, there was bias towards the 3’-end of the gene body in case of the 5’-end sequence and full-length sequence protocols. The bias towards the 3’-end indicated by high coverage at the 3’-end of the gene body (**Fig. 1c**). The tag-based sequencing of 5’ or 3’ ends methods ^[11-14]^ should have the peak coverage at either the 5’ or 3’ end of the gene with low/no coverage in the middle region of the gene body. In the second type, there was high coverage in the middle of the gene for 5’-end and 3’-end sequence protocols (**Fig. 1c**), in contrast to the full-length sequencing protocols. In the third type, there was low coverage in the middle of the gene for full-length sequence protocols. This indicated by low coverage at mid-point of 5’-3’-end of gene body (**Fig. 1d**).

The above variation (bias) and skewness in the gene body coverage among individual cells of the same dataset reflect the success of each single-cell reaction. The variation in the gene body coverage of individual cell should be considered when analyzing scRNA-seq data. The computed gene body coverage for all dataset available for download via SCPortalen database (**Supplementary Notes**).

### Variation in gene expression and gene saturation among scRNA-seq protocols

To analyze the variability in gene expression for each of the scRNA-seq protocols, we computed normalized gene expression for each dataset (**Online Methods**). The variability in gene expression among single-cells of the same cell type generated by different protocols is illustrated in (**Fig. 1e and f**) and (**Panel c and d in Supplementary Figs. 1-4; and Panel c in Supplementary Fig. 5 and 6**). The variability is more visible in the 5’ end and 3’ end sequence protocol compared to the full length sequence. We notice higher coefficient of variation over the mean (CV/mean) for the 5’-end and 3’-end sequence protocols, and lower CV/mean for the full length sequence protocols.

We examined the correlation between the same batch-matched cell type (**Supplementary Fig. 7**). The figure illustrates the variability in the mean expression of scRNA-seq protocols; the two Mouse embryonic stem cells (mESCs) dataset (E-MTAB-2600^[15]^ and E-MTAB-2805^[16]^) produced by different labs using the SMARTer protocol showed strong correlation (R^2^ = 0.85). Comparison of the mean expression of the mESCs dataset generated by SMARTer (E-MTAB-2600)^[15]^ and SUPer-Seq (GSE53386)^[17]^ showed weak correlation (R^2^ = 0.61). SMARTer has better mean expression correlation with full length sequence protocols compared to the 5’-end and 3’-end sequence protocols. The mESCs dataset (GSE46980)^[4]^ and (GSE29087)^[18]^ were 6 generated by two different versions of STRT^[11]^ protocols and showed weak correlation (R^2^ = 0.54). In general, the mean gene expression values of datasets from the same cell type (mESCs) generated by different protocols were dissimilar. Similar patterns as for (mESCs) were illustrated using mouse CD4 T-cells, mouse fibroblasts, mouse hematopoietic cell and human embryonic stem cells (hESCs) (**Supplementary Figs. 7–11**).

To investigate the variability in gene saturation for different scRNA-seq protocols, we used the Hanabi plot ^[19]^ The Hanabi plot considers the number of detected genes over the total counts. The figures (**Supplementary Figs. 12–18**) demonstrate the detection power and variability in gene saturation for different protocols. The 5’-end and 3’-end sequence protocols detected smaller numbers of genes with smaller total read counts compared to the full-length sequencing protocols.

### SkewC segregate typical and poor quality single-cells

The results from the gene body coverage analysis discriminate two classes of cells, referred to as typical single-cells and skewed single-cells, even in one dataset. A method is required to segregate typical cells and skewed cells in any scRNA-seq dataset to avoid biases and large variation. To systematically classify the two classes of the cells, we developed an algorithm (SkewC). SkewC takes as input the indexed BAM file of single-cell and the gene model (**Fig. 2**). In some of the dataset, we removed single-cells with low numbers of mapped input reads (left charts of **Fig. 3a**). The remaining single-cells had high numbers of mapped input reads. We applied the method systematically for all datasets in this study (**Fig. 3a-c**) and (**Supplementary Figs. 19–24**). When applying our method to all datasets, two distinct clusters of single-cells are visible (middle charts of **Fig. 3a-c** and middle charts of **Supplementary Figs. 19–24**).

**Figure 2:**
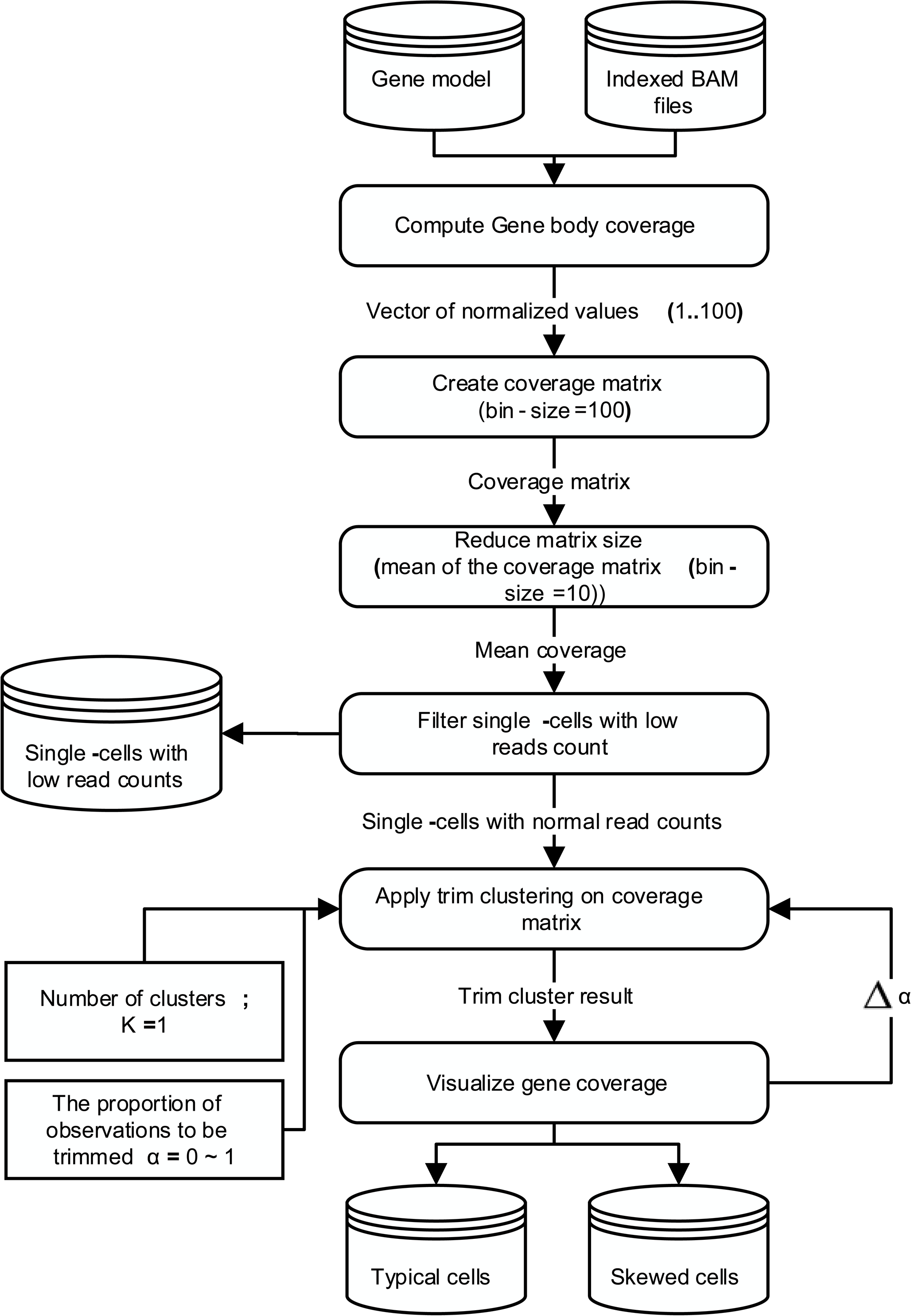
Implementation of SkewC method. Overall description and steps to implement SkewC; The figure illustrates the method to discriminate skewed cells with skewed coverage distribution.

**Figure 3:**
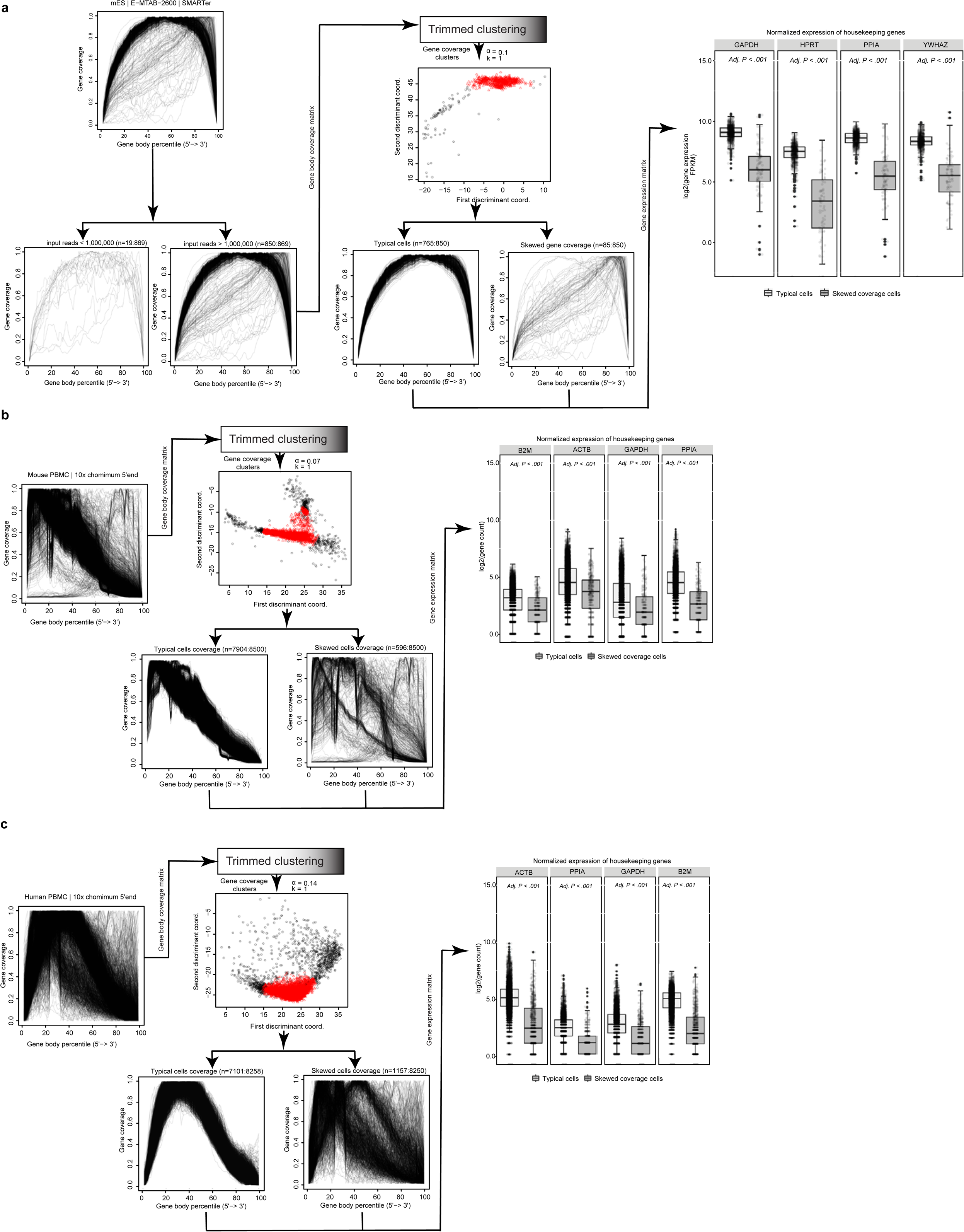
Classification of the typical and skewed coverage distribution cells. Panels (a–c) application of SkewC in 3 different datasets. Left chart filter cells with low-input reads, middle chart performs trimmed clustering on the coverage matrix of the cell with high read count. A proportion of the most outlying observations is trimmed (the skewed cells). This results into two sets of cell: typical cells with normal gene coverage and skewed cells with skewed coverage distribution. Right chart, the classification of cells was validated with the housekeeping genes (boxplot of the expression of the house keeping genes of the typical vs. skewed cells.). We applied our method to other dataset (Supplementary Figs. 19–24)

The ratio of the typical cells to skewed cells is different between different datasets. As an example, the mESCs dataset E-MTAB-2600^[15]^ has a total number of single-cells (n=869), of which single-cells with low input mapped reads (n=19), typical single-cells (n=765) and skewed single-cells (n=85). As another example, the dataset GSE98664^[20]^ with a total number of single-cells (n=364), there were no single-cells with low input mapped reads (n=0), typical cells (n=338) and skewed cells (n=26).

### Expression variation of housekeeping genes between typical and skewed cells

To investigate the difference between the typical and skewed cells in the resulting gene expression, we compared the normalized expression of the housekeeping genes (HKGs) of the typical cells versus the skewed cells (boxplot in **Fig. 3a-c** and **Supplementary Figs. 19–24**). The boxplot shows distinct differences in the variability in gene expression of the HKGs between the two classes of cells with adjusted P-values < 0.001.

### Validation of SkewC and the biological features of the skewed cells

To validate SkewC, we used the mESCs dataset GSE46980^[4]^. The authors of the dataset classify the single-cells in their experiment as good quality cells (n=47), poor quality (n=40) and dead cells (n=9), totaling n=96 single-cells. We applied SkewC on the live single-cells (n=87). The *t*-distributed stochastic neighbor embedding (*t*-SNE) ^[21]^ plot (**Fig. 4a**) shows a clear distinction of the typical and skewed cells. SkewC reduced the number of typical cells from n=47 to n=39 and increased poor quality from n=40 to n=48. This indicates that the standard QC procedures currently used in scRNA-seq analysis are inadequate to discover single-cells with poor quality.

**Figure 4:**
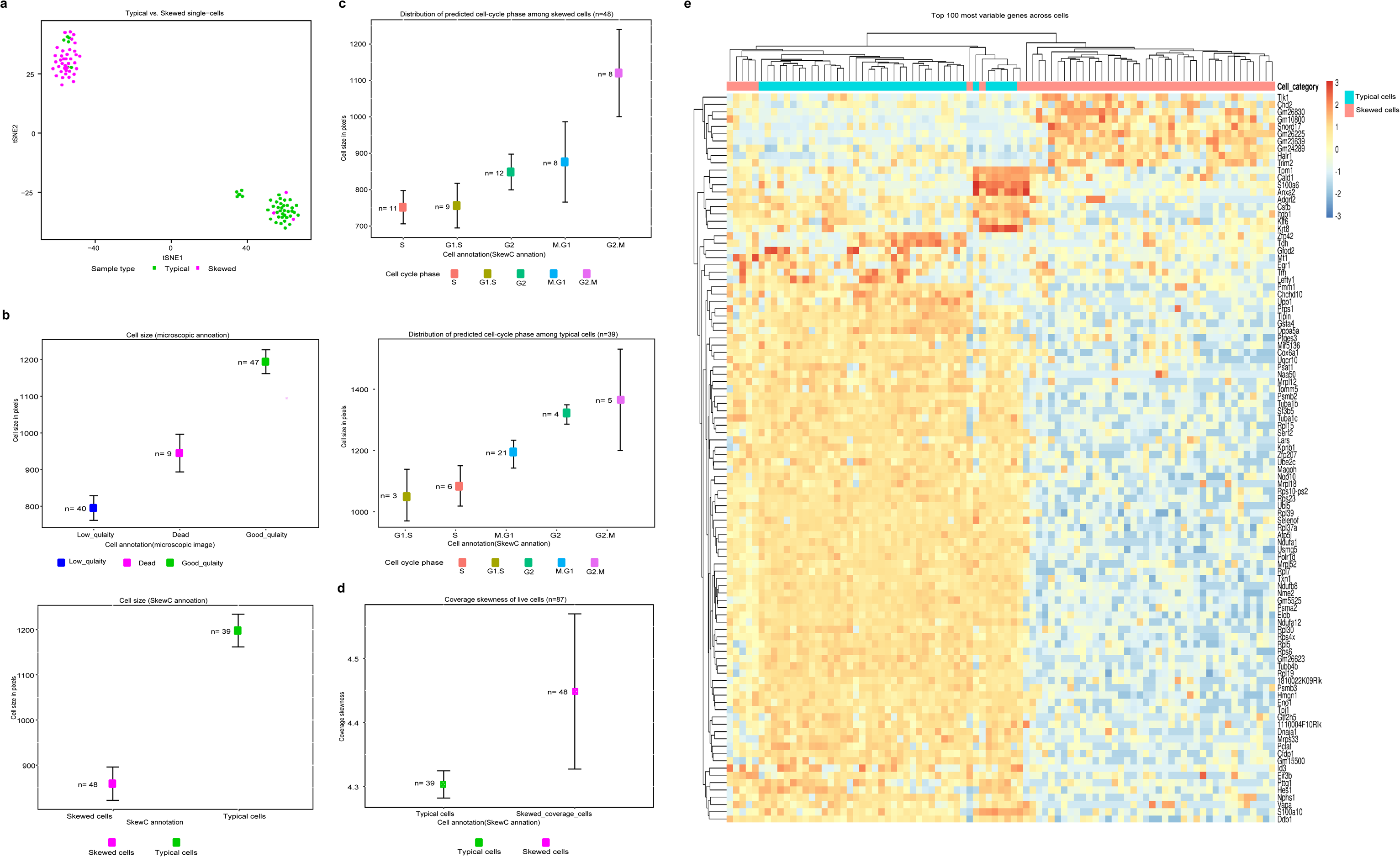
Validation of SkewC method and biological features of skewed cell using GSE46980. We used an existing experimentally validated dataset to validate the QC methods. The dataset GSE46980. (a) *t*-SNE illustrating the clustering of single-cells. (b) Line and point graph with error bars representing the standard error of the mean of the single-cell size. Upper panel shows that cell size of the single-cells as annotated by the dataset authors (dead, low quality and good quality cells). The bottom plot shows that cell size of the cells as annotated after applying SkewC method (Skewed cells and typical cells). (c), distribution of predicted cell-cycle phases among skewed cells (top) and typical cells (bottom). (d) Coverage skewness comparison between typical and skewed cells. (e) Heatmap showing the top 100 differentially expressed genes in the dataset. Columns are cell category (typical and skewed) and rows are gene names.

To investigate the biological features and meaning of the typical and skewed cells we used the microscopic image of the Fluidigm C1 chip provided by the authors in ^[4]^ and computed the cell size in pixels of all single-cells (**Supplementary Table 1**). The typical cells have larger cell size compared to the skewed cells (**Fig. 4b**, bottom plot).

An additional analysis to investigate biological difference between typical and skewed cells is the distribution of the cell-cycle phase in the dataset. We assigned the cell-cycle phase for each single-cell based on their gene expression (**Online Methods**). The majority of the typical cells were in the G2/M and M/G1 phase (n=26 of 39; bottom graph of **Fig. 4c**), suggesting that typical cells pass the G2 checkpoint (G2/M) and the spindle checkpoint (M/G1). On the other hand, the majority of the skewed cells are in G1/S, S and G2 phase (n= 32 of 48), indicating that skewed cell reside around the S phase (Synthesis Phase) but do not pass to the Mitotic phase (chromosome separation, top panel of **Fig. 4c**).

We investigated the differences in gene coverage distribution (skewness) between the typical and skewed cells (**Fig. 4d**). The skewed cell possess high coverage skewness compared to the typical cells.

Finally, we performed differential gene expression analysis between the typical and skewed cells. The clustering of the top 100 most variable genes across cells illustrated in the heatmap (**Fig. 4e**) with typical and skewed cells are clustered separately based on the gene expression of the top 100 genes. We performed gene set enrichment analysis (GSEA) on of the top 100 genes, and found that the KRAS signaling DN pathway was enriched in the typical cells (p-value < 0.001). The set of genes up-regulated by KRAS play roles in cell signaling^[22]^.

### Effect of skewed cells on downstream analysis

Since we noticed a great variability in the mean expression of the HKGs between typical and skewed cells, we investigated the effect of skewed cells on the downstream analysis of the scRNA-seq experiments. The *t*-SNE ^[21]^, is a common dimension reduction technique in scRNA-seq analysis. *t*-SNE is usually performed after read count normalization ^[23]^. We analyzed the impact of filtering skewed cells on *t*-SNE implementation using mESCs dataset (**Fig. 5**). In the first dataset E-MTAB-2600^[15]^ generated by SMARTer protocol (**Fig. 5a**), the top panel shows the *t*-SNE plot of all single-cells clustered and colored by the growth factors used in the experiment, some of the single-cells are misplaced in the wrong cluster (mis-cluster). The *t*-SNE plot in the middle panel of (**Fig. 5a**), cluster and color the single-cells based on the classification of typical and skewed cells. The majority of the skewed cells clustered together with few exceptions. The *t*-SNE plot in the bottom panel of (**Fig. 5a**) shows replotting the *t*-SNE after filtering the skewed cells. The plotting of the typical cells only show distinct clustering of the cells based on the growth factors compared the *t*-SNE before the filtering of the skewed cells. This shows the impact of the skewed cells on the clustering of the single-cells.

**Figure 5:**
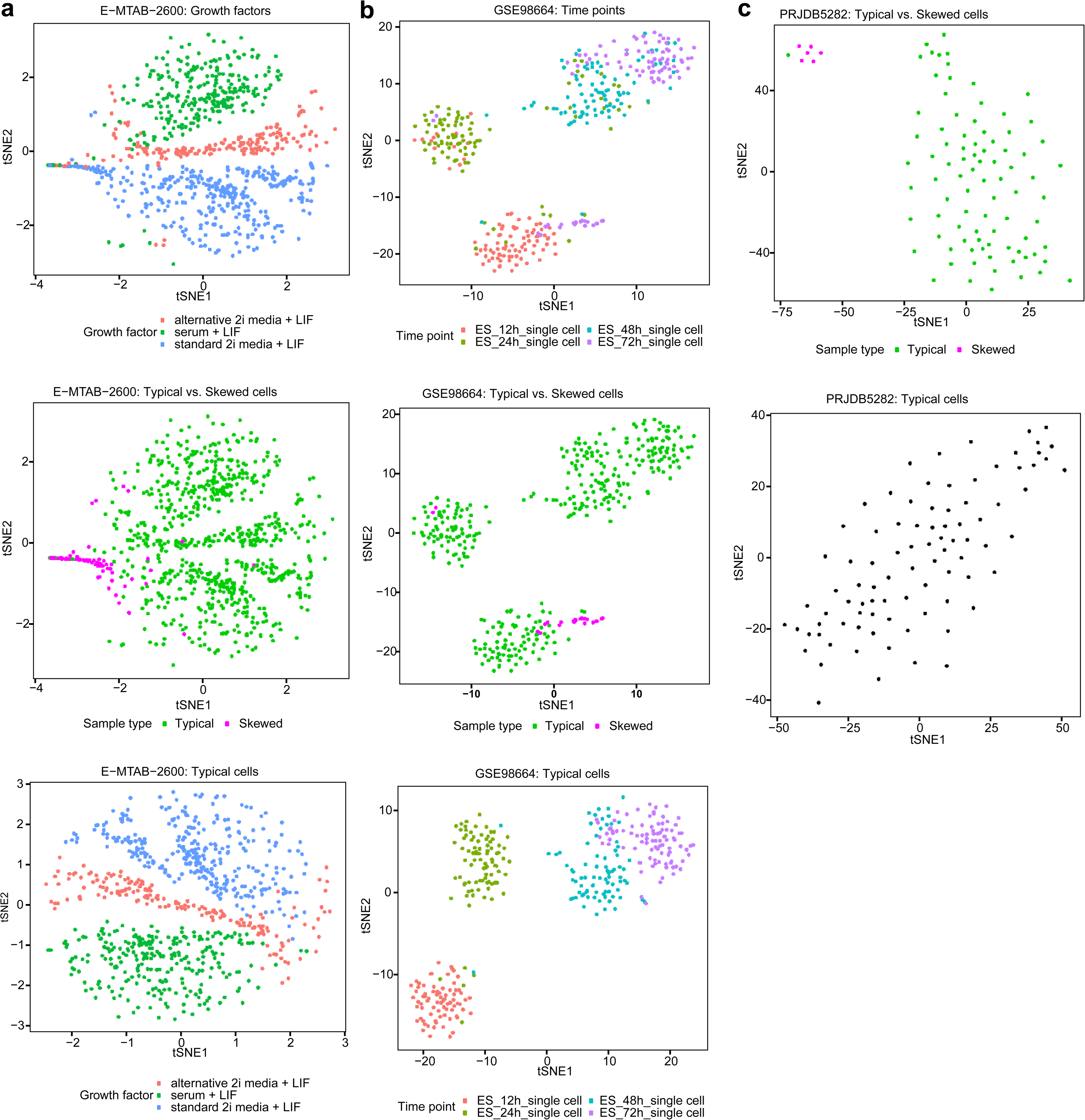
Effect skewed cells on downstream analysis. To evaluate the effect of the skewed cells when performing downstream analysis, t-SNE generated for several datasets. (a) mESCs single-cell treated with three types of growth factors. On the top is the t-SNE before the classification and colored by the growth factor used, in the middle a t-SNE of the skewed and typical cells, on the bottom t-SNE after removing the skewed cells. (b) mESCs with four development time-points. Top t-SNE of the dataset before the classification. The cluster in the bottom show the majority of the cells at 12 h, with few cell from 72 h. In the middle t-SNE we observed that the skewed cells are cells from 72h. The bottom t-SNE show better clustering of the single-cell per development time-point after filtering the skewed cells. (c) t-SNE perfectly discriminate typical and skewed cells.

The dataset GSE98664^[20]^ is a time-course analysis of mESCs development generated by RamDA-seq protocol (**Fig. 5b**). The top panel illustrates *t*-SNE with four clusters (four time-points). Each of the four clusters contains single-cells that do not belong to the same time-point (mis-clustered). The wrongly clustered single-cells are in fact skewed cells that stopped development but were mis-annotated by as developing cells. The middle *t*-SNE plot shows the clustering of the same dataset based on typical and skewed cells and the appearance of the skewed cells in two clusters (ES_12h & ES_24h). The bottom *t*-SNE illustrates re-clustering of the dataset after filtering the skewed cells; the plot shows clear improvement of the clustering result. Compared to the top *t*-SNE plot, the final *t*-SNE after filtering of the skewed cell removed the group of single-cells of time-point ES_72h from ES_12h and the ES_24h cells were removed from the cluster from ES_48h.

In the single-cell dataset PRJDB5282^[13]^, generated by C1 CAGE protocol (**Fig. 5c**), the skewed cells clustered separately on *t*-SNE plot (top of **Fig. 5c**). After filtering the skewed cells, *t*-SNE shows one cluster consisting of typical cells.

All the above examples demonstrate the strong impact of skewed cells on the clustering results. The identification and filtering of the skewed cells is important to consider in any downstream analysis.

### Ratio of intergenic expression

In scRNA-seq protocols, cDNA is obtained from the reverse transcription of RNA. This step is followed by amplification of cDNA by PCR or in vitro transcription before sequencing^[24]^. The amplification step is required due to the small amount of RNA found in an individual cell, and the workflow is prone to losses or biases^[25]^. To investigate the possibility of such problems resulting, e.g., from genomic DNA contamination, we quantify the ratio of intergenic expression for each cell (**Online Methods**). As a control, we considered matched cell type bulk RNA-Seq dataset from ENCODE^[26]^ (PloyRNA-Seq and Total RNA-Seq). As an example (**Fig. 6a**), the dataset GSE68981^[27]^ from mouse hematopoietic stem cells (HSCs) was analyzed with single-cell (C1-single-cell mRNA-Seq protocol) and bulk RNA-Seq used as control. As another example, we compared the human datasets GSE75748^[28]^ (**Fig. 6b**), from human embryonic stem cells (analyzed with single-cell SMARTer protocol) and bulk RNA-Seq. As demonstrated in the above examples, the ratio of intergenic expression is high in the scRNA-seq data compared to the bulk RNA-Seq.

**Figure 6:**
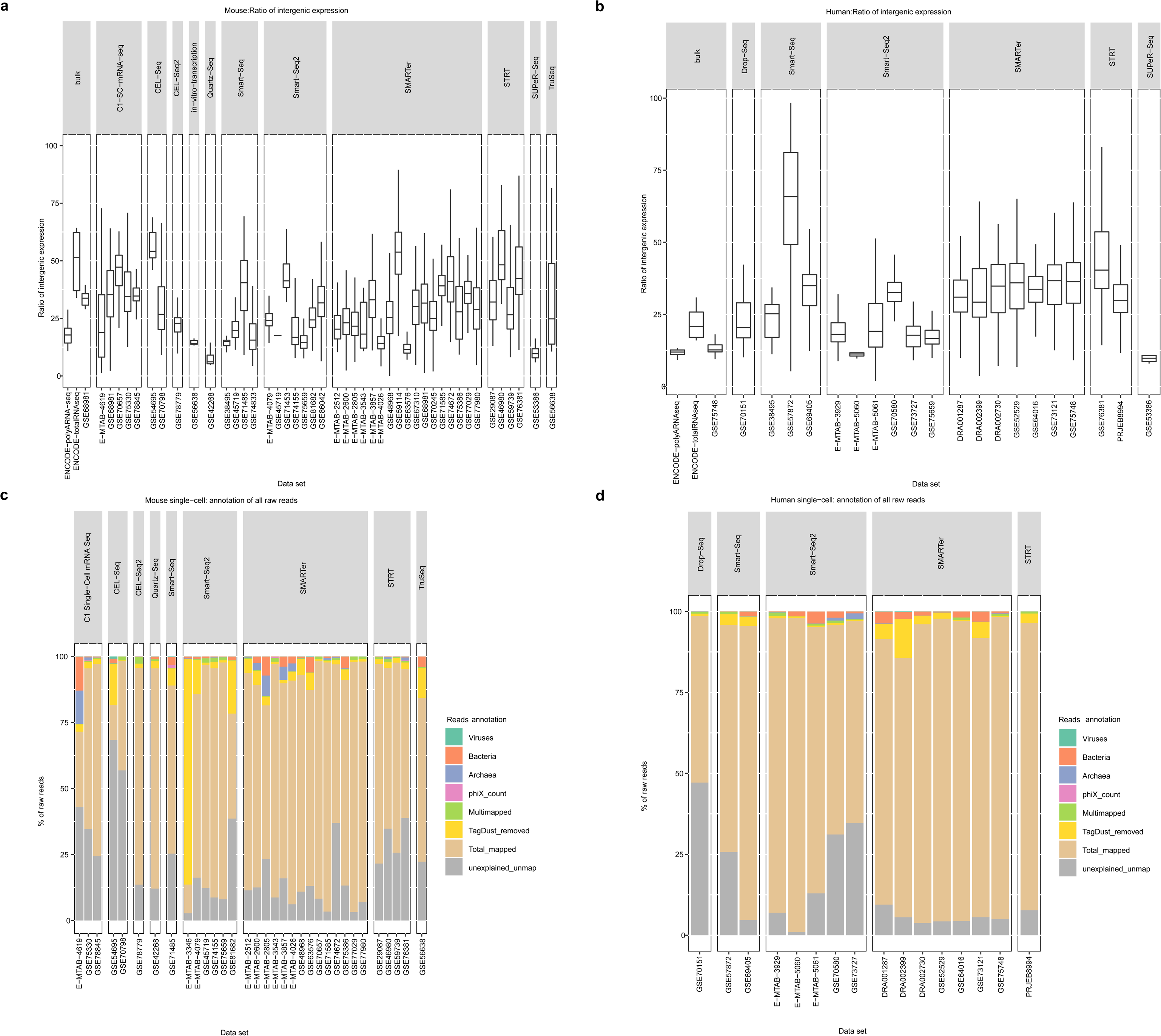
Ratio of intergenic expression and annotation of the reads. (a–b) High level of intergenic expression in scRNA-seq (a) mouse and (b) human data compared to bulk. Box plots are grouped by protocol name. (c–d) Classification of reads annotations per dataset for (c) mouse and (d) human data. Mapped reads are separated between uniquely mapped and multimapped on the reference genome. Unmapped reads are annotated in categories according to the explanation of unmapping, namely tagdust filtered, positive sequencing control (phiX), contamination by a foreign organism (archea, bacteria, virus). Unmapped reads that couldn’t be explained are presented in gray.

Our analysis suggested that scRNA-seq data prone to high level of intergenic expression compared to bulk RNA-Seq. One possibility for such high read counts from intergenic regions is amplification of genomic DNA, as such signals are not observed in bulk RNA-seq data from same cell types. Our results suggest that single-cell dataset should be evaluated for intergenic expression, possibly originating from genomic DNA amplification.

### Annotation and classification of the sequence reads

One potential source of problems with scRNA-seq is microbial contamination (bacteria, archaea and viruses), we developed a workflow to annotate and classify unmapped sequence reads from scRNA-seq experiments (**Online Methods**). Sequence reads are annotated on three categories, single-mapped, multi-mapped, or unmapped on the reference genome. The majority of the sequence reads are annotated as mapped reads (**Fig. 6c-d**) (wheat color) with one exception of the dataset E-MTAB-3346^[29]^ (mouse thyme epithelial cells (mTECs) generated by Smart-Seq2 protocol) (**Fig. 6c**). The unmapped reads are annotated as positive sequencing control (phiX), or as reads belonging to unexpected organisms (bacteria, archaea, virus) (**Fig. 6 c-d**). The dataset PRJEB8994^[1]^ of gene expression during the first three days of human development (**Fig. 6d**) is the only dataset that doesn’t contain any reads annotated as unexpected organisms (bacteria, archaea, virus), possibly because the PRJEB8994 experiment was conducted in a sterile clinical environment.

A set of unmapped sequence reads was annotated as unexplained-unmapped (**Fig. 6 c-d**) (gray color). These are reads that failed to find any match in the Kraken database. Notable, the two datasets GSE54695^[30]^ and GSE70798^[31]^ generated by CEL-Seq contain high numbers (> 50% of the raw reads) of unexplained-unmapped reads. A common source of the unmapped reads in scRNA-seq experiments might arise from the sequence linkers. Specific sequencing linkers are usually added to cDNA during library construction. The raw data files for the sequence reads analysis and annotation are available for exploration through interactive user interface and for bulk download via the online database as described in the (**supplementary information**) Mycoplasma contamination of cell cultures in the laboratory is common and special detection kits are available to ensure that the cell culture is free of such contamination. In bulk RNA-seq, it was reported that several large scale RNA-seq projects generate 9-20% of reads that do not map to the human reference genome^[32]^.

## Discussion

Advances in scRNA-seq have already impacted biology and medicine and will increasingly do so. It has enabled investigation of transcriptomic variation between individual cells, thereby enabling the discovery of new cell types, analyses of cellular response to stimulation, analyses of the nature and dynamics of cell differentiation and reprogramming, and study of transcriptional stochasticity ^[25]^. In spite of the technical advances, several challenges still remain and need to be understood for improved interpretation of the data. There is high variability in the performance of scRNA-seq protocols in terms of coverage, accuracy and specificity, impacting the quality of data generated by different scRNA-seq protocols^[33, 34]^. The variability among scRNA-seq protocols and the quality of the scRNA-seq dataset might also impact global efforts to map transcription in human and mouse cells^[35, 36]^.

Our analysis identified wide differences in patterns of gene coverage generated from same cell types by different scRNA-seq protocols. Based on the analysis of the gene body coverage, we identified two classes of single-cells in any dataset: cells with method-specific (prototypical) distribution of gene body coverage, and cells with a skewed distribution. Each of the scRNA-seq protocols yields different gene body coverage. A skewed distribution with excess observed 3’-end bias could be attributed to technical or biological processes during any of the experimental steps, such as reaction failure or cell death, triggering mRNA degradation. During embryo development, skewed distribution coverage suggested induction of maternal transcript degradation at early genome activation in the earliest developmental stages^[37]^, as maternal transcript degradation takes place after the fertilization till day 3^[1, 38]^.

SkewC is based on the observation of the two distribution patterns in the gene body coverage of scRNA-seq protocols and the identification of the typical and skewed cells. The skewed cells might result from a technical failure of the reaction, cell death, or a biological process as in the embryo development. The skewed live cells might potentially be quiescent satellite cells, a form of (**G_0_** cell-cycle phase), cell enter **G_0_** phase (resting) due to different environmental factors. The common feature between the identified skewed cells and the quiescent satellite cells is the low RNA content^[39, 40]^.

We can find the impact of gene body coverage skewness in many aspects. In the expression of housekeeping genes, we identified significant differences between typical and skewed cells. Regarding the cell size and cell-cycle of typical and skewed cells, typical cells show strong correlation between cell size and cell cycle phase, in contract to skewed cells. The differential gene expression analysis demonstrated that skewed cells show different expression profile of the top expressed genes compared to the typical cells. We further assessed the impact of filtering out the skewed cells from downstream analysis (clustering and differential gene expression analysis), and found that exclusion of skewed cells drastically changes their clustering results. Wrong clustering may lead to false discovery resulting from the skewed cells. From these results, we recommend that skewed cells should be identified and excluded from any downstream analyses of scRNA-seq experiments.

The dataset-to-dataset similarity analyses in terms of gene expression profiles rather expectedly show weak expression correlation between datasets generated by different protocols. Our results show strong variation in sequence depth, detection power and gene saturation revealed by different scRNA-seq protocols. This result demonstrated challenges for the current efforts to computationally integrate heterogeneous scRNA-seq datasets generated by different protocols and labs^[41-43]^.

We estimated the intergenic expression in scRNA-seq, and observed high level of intergenic expression in single-cells compared to the control bulk dataset. This might be potentially caused by scRNA-seq contamination with genomic DNA reads, but other alternatives to explain this observation might also exist. Finally, we annotated and classify unmapped reads in order to find the source of contamination in scRNA-seq experiments. This result suggests the possible contamination of virus or bacterial sequences in the scRNA-seq results.

In conclusion, our results demonstrated that the presence of skewed cells influence the data analysis and interpretation and therefor a QC method to segregate typical cells and skewed cells in scRNA-seq dataset should be standard procedure in any scRNA-seq experiment. Indeed, SkewC method described here, is novel and can easily be integrated in scRNA-seq data analysis workflows.

## Online Methods

### Study design

Based on the objective of the scRNA-seq experiment, the protocols are divided in two categories; full-length sequence profiling or transcript end-tagging (5’ or 3’). In full-length sequence protocols (SMARTer, Smart-Seq, SUPer-Seq, RamDA-seq, etc.) the sequence reads cover the entire gene body (5’to 3’-end) and quantify gene and transcript isoforms. The end-tagging based sequencing protocols (C1 CAGE, CEL-Seq, CEL-Seq2, STRT, 10x Chromium Single Cell 3’ –end etc.) target one end of the transcript (5’ -end or 3’ -end) and are used to identify promoters (5’ tagging) or give an estimate of transcript abundance. We compare and analyses scRNA-seq protocols using approaches different from the published studies ^[33, 34]^. In particular, we evaluate the capability and power of each protocol in terms of the full-length transcript coverage, variability in sequence depth, expression variation, ratio of intergenic expression and analysis of the unmapped sequence reads to the reference genome (**Supplementary Fig. 25**). We analyzed 29 datasets produced with 15 different protocols representing commonly used scRNA-seq methods of the above two categories. Based on the target read capture strategy of the scRNA-seq method, 5 datasets measured gene expression at the 5’-end of the transcript (STRT, C1 CAGE and 10x Chromium 5’-end), 20 datasets measured gene expression of the full-length transcript (SMARTer, Smart-Seq, RamDA-seq, SUPer-Seq, Quartz-Seq, C1 single-cell mRNA-Seq, Smart-Seq2, TruSeq and Drop-Seq), and 4 studies measured gene expression at the 3’-end of the transcript (CEL-Seq, CEL-Seq2, and 10x Chromium 3’-end).

### Study dataset

To perform fair comparison among different protocols and to make the results comparable, we designed the datasets as batch-matched cells and unmatched cells (**Supplementary Table 2**). For the mouse batch-matched cells, we analyzed mouse embryonic stem cells (mESCs), CD4 T cells, fibroblasts and hematopoietic cells. For human batch-matched cells, we analyzed human embryonic stem cells (hESCs). The mouse unmatched cells were adipocytes and PBMC cells, and the human unmatched cells were MCF10A cells, PBMC cells and HEK 293 and 3T3 cells (**Supplementary Table 2**).

The analyzed datasets covered tissues, primary cells and cell lines. We generated a dataset for human MCF10A cells by the 10x Chromium 5’ -end protocol. We reanalyzed published human and mouse scRNA-seq from International Nucleotide Sequence Database Collaboration (INSDC) dataset. Finally, we utilized 10x Genomics data resource and reanalyzed published scRNA-seq dataset for human and mouse from 10x Genomics data portal. The three types of the datasets are described below.

### 10x Genomics Chromium experiment of human MCF10A cells

10x Genomic Chromium dataset was generated from the MCF10A cells (ATCC). The cells were grown in DMEM/F12(1:1) as described in ^[44]^. RNA-Seq library was prepared using 10x Chromium Single Cell 3’ –end Reagent Kits User Guide (v2 Chemistry). Libraries were sequenced using paired-end sequencing (26bp Read 1 and 98bp Read 2) with a single sample index (8bp) on the Illumina HiSeq 2500. Raw reads and processed data are deposited under GEO GSE143607.

### International Nucleotide Sequence Database Collaboration (INSDC) dataset

The method for collecting and processing the raw read scRNA-seq from the public databases were illustrated in ^[45]^ and listed in **Supplementary Table 3**[1, 4, 12, 13, 15-18, 20, 27, 28, 30, 46-59]. In brief, published scRNA-seq were collected by searching PubMed for human and mouse scRNA-seq articles. Our strategy was to include different types of cells generated by different technology platforms. We retrieved study accession number(s) of the original data deposited to International Nucleotide Sequence Database Collaboration (INSDC). The study accession numbers were used to retrieve sequence read files and metadata files from INSDC sites (DDBJ, EMBL-EBI, NCBI). To obtain FASTQ files, we implemented an automated program using the NCBI SRA Toolkit^[60]^. Metadata about each dataset were collected as well. This metadata contains information about the cell type, protocol, sequence platform, single-cell isolation techniques, etc. We implemented automated script to retrieve dataset metadata utilizing The Entrez Programming Utilities (E-utilities) from NCBI^[61]^.

### 10x Genomics dataset

We downloaded two datasets from 10x Genomics portal (https://support.10xgenomics.com/single-cell-vdj/datasets). The Human PBMCs of a healthy donor - 5’ –end gene expression and cell surface protein (8,258 single-cell) BAM files were downloaded from http://cf.10xgenomics.com/samples/cell-vdj/3.0.0/vdj_v1_hs_pbmc2_5gex_protein/vdj_v1_hs_pbmc2_5gex_protein_web_summary.html The Mouse PBMCs from C57BL/6 mice - 5’ gene expression (8,500 single-cell) BAM files were downloaded from http://cf.10xgenomics.com/samples/cell-vdj/3.0.0/vdj_v1_mm_c57bl6_pbmc_5gex/vdj_v1_mm_c57bl6_pbmc_5gex_web_summary.htm l

### Data processing of the raw sequence reads

For the raw sequence data downloaded from INSDC, we run basic QC procedures to obtain quality assessment metrics of the raw sequence reads. The QC procedures includes testing of all FASTQ files with FastQC tool [http://www.bioinformatics.babraham.ac.uk/projects/fastqc/] to identify any quality issues. Additional QC procedures of the raw reads includes count of raw tags. The raw sequence reads were aligned to a recent reference genome build (GRCh38 (human) or GRCm38 (mouse) genome assembly). We used STAR software (version 2.5.1b)^[62]^ with default settings and GENCODE gene annotations in the release v24 for human and the release vM9 for mouse for all dataset but not for (GSE98664) in which we used GENCODE vM22. Aligned reads in BAM file format together with the log files generated by STAR were used to obtain quality assessment metrics (total read count, number of uniquely mapped reads and assigned reads (mapped reads assigned to gene)). The mapping ratio, counts of mapped reads, unmapped and multi-mapped reads were summarized using SAMtools software^[63]^. For MCF10A dataset, we used Cell Ranger version 2.1.1 for data processing. The datasets from 10x Genomics were processed with Cell Ranger version 3.1.0. The implemented workflow and scripts are publicly available (see **Supplementary Notes**)

### Reads count summarization and expression normalization

To obtain expression matrices, we quantified gene expression counts using featureCounts (in the Subread package Version 1.5.0-p1) ^[64]^. The gene expression counts were normalized into tags 19 per million reads (TPM) for 5’-end methods and fragments per kilobase million (FPKM) for other methods to generate a gene expression table for each study, according to the following standard formula.

TPM (gene-level expression) = mapped reads assigned to each gene) * 1,000,000 / mapped reads FPKM (gene-level expression) = mapped reads assigned to each gene) * 1000 / (gene length (bp)) * 1,000,000 / mapped reads. For the 10x Genomics dataset we used the computed feature-barcode matrices MEX provided by Cell Ranger software.

### Computation and visualization of the gene body coverage

To compute the gene body coverage for dataset, we used the geneBody_coverage.py program in RSeQC ^[65]^. The program was used to check if reads coverage was uniform and if there was any 5’/3’ end bias. The input for the program is indexed BAM files and gene model in BED format. Gene models were downloaded from http://rseqc.sourceforge.net/#download-gene-models-update-on-08-07-2014. The gene model BED files for human hg38_Gencode_V28.bed.gz and for mouse mm10_Gencode_VM18.bed.gz were preprocessed to filter ribosomal RNA and transfer RNA.

For 10x Genomics dataset, we postprocessed the barcoded BAM file to split it into individual BAM files using the cell barcodes, which gave us one BAM file per each single-cell.

Since processing of the gene body coverage is a time-consuming task even in a high performance computer environment, datasets with large numbers of single-cells were split to smaller chunks of datasets to run small jobs in parallel. The result of the program is a vector of values consisting of normalized coverages within its gene body from the 5’-end to the 3’-end binned to 0-100 (positions). The value for each position range from (0-1), where 0 indicate no coverage and 1 indicate full coverage at the position on the gene body. The vector of the normalized values was postprocessed in several steps for visualization and study the normality and skewness of the coverage (**Fig. 2**).

### SkewC implementation

SkewC is implemented in R and the code is available with test data in GitHub (see **Data Availability**). As explained in (**Fig. 2**). The following are the main steps implemented in SkewC:

1. Filter cells with low-input reads from the dataset (Input raw counts table)
2. Perform trimmed clustering using tclust R function on the coverage matrix (Input processed gene body coverage matrix)
  a. Set number of cluster ***K***= 1 (constant)
  b. Instead of trying to fit noisy data, a proportion of the most outlying observations is trimmed. This proportion (α), is variable
3. Visualize gene coverage of the result of trimmed clustering. This will result in a set of single-cell with typical gene coverage and a set of single-cell with skewed gene coverage. Based on the visualization of the gene body coverage of the trimmed clustering, α can be changed and user re-run step#2 and #3 several times (iteration process).

### Computation of intergenic expression

To quantify the ratio of intergenic expression (e.g. due to the PCR amplifications of the starting material of genomic DNA), we modelled the following formula:

The ratio of intergenic expression = (((total number of mapped reads) - (the total number of reads that are assigned to a gene feature in GENCODE annotation)) / ((total number of mapped reads))) * 100.

The ratio of intergenic expression was computed for each single-cell and summarized per the dataset.

### Analysis and annotation of the unmapped reads

In our pipeline for aligning raw reads to the reference genome, we kept log files and unmapped reads. We investigated the source and ration of the unmapped reads from each FASTQ file. Our workflow to analyze the unmapped reads started with filtering of ribosomal RNA and artificial reads from the BAM file of the unmapped reads using TagDust tool^[66]^. Next we filtered multi-mapped reads to get true unmapped reads and finally, we performed blast search on the most frequent unmapped reads ^[67]^. The remaining reads were screened for microbial contamination sequences. To screen for such organisms, we utilized metagenomics tools: sequana^[68]^ and Kraken ^[69]^. The Kraken tool provided 8GB database of complete bacterial, archaeal and viral genomes (https://ccb.jhu.edu/software/kraken/dl/minikraken_20171019_8GB.tgz).

### Cell size estimation of the dataset GSE46980

The dataset GSE46980 of mESCss ^[4]^ was generated by the STRT protocol and provided full annotation of quality status of the single-cells (n=96). The authors classify each single-cell as either dead (depleted before cell capture by the flow-cell) or live cell. The live cells were further classified as either low quality cells or good quality cells (see ^[4]^ for details on how the annotation was performed). We used this dataset to compare our QC method of typical and skewed cells. Additionally, the dataset provided a microscopic image of the Fluidigm C1 chip. In the microscopic image, each Fluidigm C1 chip (a 96-well plate) was imaged after cell capture and a grid of thumbnails was generated for each chip. To verify some of the morphological phenotypes of the typical and skewed cells, we estimated morphological properties of the cells based on the microscopic image, cell size, areas, circularity, skewness roundness and solidity that were calculated using the ImageJ tool ^[70]^.

### Cell-cycle phase prediction

To further evaluate some of the biological phenotypes of the typical and skewed single-cells, we predicted the cell-cycle phase, the cell-cycle phase predicted computationally based on the expression profile of the single-cell^[71]^. We obtained the predefined human cell-cycle marker set provided in ^[72]^. As for the mouse cell-cycle markers, the orthologous mouse genes of the human cell-cycle gene markers were obtained (**Supplementary Table 4**). The cell-cycle phase predictor ^[71]^ assign any of (S, G1/S, G2, M/G2, G2/M) phase to each single-cell.

### Statistical tests, boxplots and plotting tools

Unless otherwise indicated, all p-values were obtained with two-sided t-test. In all boxplots, center lines indicate median values, box heights indicate the inter-quartile range of data. The ggplot2 library from R software version 3.5.1 (2018-07-02) was used for plotting of all plots and figures.

### Data availability

The raw read data listed in (**Supplementary Table 2**) are available from INSDC sites. We developed SCPortalen, a single-cell database in which we deposited all results from this study at http://single-cell.clst.riken.jp/. A full description and use guide provided in the (**Supplementary information – Online Resources**) The scRNA-seq dataset for human MCF10A cells generated in this study have been deposited in the NCBI GEO under accession GSE143607.

The R code for SkewC and all scripts used to process dataset from International Nucleotide Sequence Database Collaboration (INSDC) are available from GitHub https://github.com/LSBDT

## Acknowledgements

This work was supported by research grants for the RIKEN Center for Life Science Technologies, RIKEN Center for Integrative Medical Sciences and RIKEN Open Life Science Platform project from MEXT, Japan. SK and JK were supported in part by Knut and Alice Wallenberg Foundation (KAW2015.0096) (Sweden), Swedish Research Council, Jane and Aatos Erkko Foundation (Finland), and Sigrid Jusélius Foundation (Finland). This work was initiated when JK was a Japan Society for the Promotion of Science Fellow at RIKEN Center for Integrative Medical Sciences, Yokohama.

## Supplementary Table Legends

**Supplementary Table 1: Analysis of the microscopic image for the dataset GSE46982**

**Supplementary Table 2: Sample design and tested single-cell RNA sequence protocols**

**Supplementary Table 3: List of the dataset used in the analysis**

**Supplementary Table 4: List of the cell cycle marker genes for mouse.**

## Supplementary Figure Legends

**Supplementary Figure 1: Gene coverage skewness and variation in expression among single-cell RNA-Seq protocols using mouse CD4 T cells.**

4 datasets from mouse CD4 T cells. **(a)** Distribution of the mapped reads (tags) across the genes. Each panel shows gene body coverage percentile per dataset. The x-axis represents the gene body from 5’ end to 3’ end scaled from 0-100, and the y-axis gene coverage (0-1). Each line represents a single cell. **(b)** Mean calculated for bin size = 10. **(c)** The variability in gene expression plot. The X-axis is the mean of the normalized gene expression (FPKM), Y-Axis is the coefficient of variation. CV/mean correlates the sequence depths and variability in gene expression among different protocols. **(d)** Grouping of the smooth lines from **(c)**.

**Supplementary Figure 2: Gene coverage skewness and variation in expression among single-cell RNA-Seq protocols using mouse fibroblast cells**

4 datasets from mouse fibroblast.(a) Distribution of the mapped reads (tags) across the genes. Each panel shows gene body coverage percentile per dataset. The x-axis represents the gene body from 5’ end to 3’ end scaled from 0-100, and the y-axis gene coverage (0-1). Each line represents a single cell. (b) Mean calculated for bin size = 10. (c) The variability in gene expression plot. The X-axis is the mean of the normalized gene expression (FPKM), Y-Axis is the coefficient of variation. CV/mean correlates the sequence depths and variability in gene expression among different protocols. (d) Grouping of the smooth lines from (c).

**Supplementary Figure 3: Gene coverage skewness and variation in expression among single-cell RNA-Seq protocols using mouse hematopoietic stem cells (HSCs)**

3 datasets from mouse hematopoietic stem cells (HSCs). (a) Distribution of the mapped reads (tags) across the genes. Each panel shows gene body coverage percentile per dataset. The x-axis represents the gene body from 5’ end to 3’ end scaled from 0-100, and the y-axis gene coverage (0-1). Each line represents a single cell. (b) Mean calculated for bin size = 10. (c) The variability in gene expression plot. The X-axis is the mean of the normalized gene expression (FPKM), Y-Axis is the coefficient of variation. CV/mean correlates the sequence depths and variability in gene expression among different protocols. (d) Grouping of the smooth lines from (c).

**Supplementary Figure 4: Gene coverage skewness and variation in expression among single-cell RNA-Seq protocols using human embryonic stem cells (hESCs).**

3 datasets from mouse human embryo cells. (a) Distribution of the mapped reads (tags) across the genes. Each panel shows gene body coverage percentile per dataset. The x-axis represents the gene body from 5’ end to 3’ end scaled from 0-100, and the y-axis gene coverage (0-1). Each line represents a single cell. (b) Mean calculated for bin size = 10. (c) The variability in gene expression plot. The X-axis is the mean of the normalized gene expression (FPKM), Y-Axis is the coefficient of variation. CV/mean correlates the sequence depths and variability in gene expression among different protocols. (d) Grouping of the smooth lines from (c).

**Supplementary Figure 5: Gene coverage skewness and variation in expression among single-cell RNA-Seq protocols using mouse un-matched cells.**

2 mouse dataset, the GSE56638 from Adipocyte tissue generated by Truseq and PBMC generated by 10x chromium 5’end. (a) Distribution of the mapped reads (tags) across the genes. Each panel shows gene body coverage percentile per dataset. The x-axis represents the gene body from 5’ end to 3’ end scaled from 0-100, and the y-axis gene coverage (0-1). Each line represents a single cell. (b) Mean calculated for bin size = 10. (c) The variability in gene expression plot. The X-axis is the mean of the normalized gene expression (FPKM), Y-Axis is the coefficient of variation. CV/mean correlates the sequence depths and variability in gene expression among different protocols.

**Supplementary Figure 6: Gene coverage skewness and variation in expression among single-cell RNA-Seq protocols using human un-matched cells.**

2 human dataset, the GSE70151 from HEK&3T3 cell line generated by Drop-seq, GSE143607 Human MCF10A cell line generated by 10x chromium 3’end and PBMC generated by 10x chromium 5’end. (a) Distribution of the mapped reads (tags) across the genes. Each panel shows gene body coverage percentile per dataset. The x-axis represents the gene body from 5’ end to 3’ end scaled from 0-100, and the y-axis gene coverage (0-1). Each line represents a single cell. (b) Mean calculated for bin size = 10. (c) The variability in gene expression plot. The X-axis is the mean of the normalized gene expression (FPKM), Y-Axis is the coefficient of variation. CV/mean correlates the sequence depths and variability in gene expression among different protocols.

**Supplementary Figure 7: Dataset-to-dataset similarity of the mean expression using mouse embryonic stem cells (mESCs)**

Scatterplot of the mean value of the gene expression (FPKM) to compare the variability of gene expression in mESCs dataset. Each thumbnail illustrates two datasets generated from same cell type. The dataset is either generated by same or different protocols, and some cases shows dataset generated by same protocols from different labs. The figure illustrated that gene expression of dataset from cell type, generated by different protocols are dissimilar in the average expression.

**Supplementary Figure 8: Dataset-to-dataset similarity of the mean expression using mouse CD4 T cells.**

Scatterplot of the mean value of the gene expression (FPKM) to compare the variability of gene expression in mouse CD4 T cells dataset. Each thumbnail illustrates two datasets generated from same cell type. The dataset are either generated by same or different protocols, and some cases shows dataset generated by same protocols from different labs.

**Supplementary Figure 9: Dataset-to-dataset similarity of the mean expression using mouse fibroblast cells.**

Scatterplot of the mean value of the gene expression (FPKM) to compare the variability of gene expression in mouse fibroblast cells dataset. Each thumbnail illustrates two datasets generated from same cell type. The dataset are either generated by same or different protocols, and some cases shows dataset generated by same protocols from different labs.

**Supplementary Figure 10: Dataset-to-dataset similarity of the mean expression using mouse hematopoietic stem cells (HSCs).**

Scatterplot of the mean value of the gene expression (FPKM) to compare the variability of gene expression in mouse hematopoietic cells dataset. Each thumbnail illustrates two datasets generated from same cell type. The dataset are either generated by same or different protocols, and some cases shows dataset generated by same protocols from different labs.

**Supplementary Figure 11: Dataset-to-dataset similarity of the mean expression using Human embryonic stem cells (hESCs)**

Scatterplot of the mean value of the gene expression (FPKM) to compare the variability of gene expression in human embryo cells dataset. Each thumbnail illustrates two datasets generated from same cell type. The dataset is either generated by same or different protocols, and some cases shows dataset generated by same protocols from different labs.

**Supplementary Figure 12: Hanabi plot Gene saturation using mouse embryonic stem cells (mESCs).**

The x-axis shows total counts, and y-axis shows the number of detected genes in 9 mouse ES cell dataset.

**Supplementary Figure 13: Hanabi plot Gene saturation using mouse CD4 T cells.**

The x-axis shows total counts, and y-axis shows the number of detected genes in 4 mouse CD4 T cells dataset.

**Supplementary Figure 14: Hanabi plot Gene saturation using mouse fibroblast cells.**

The x-axis shows total counts, and y-axis shows the number of detected genes in 4 mouse fibroblast cells dataset.

**Supplementary Figure 15: Hanabi plot Gene saturation using mouse hematopoietic stem cells (HSCs)**

The x-axis shows total counts, and y-axis shows the number of detected genes in 3 mouse hematopoietic cells dataset.

**Supplementary Figure 16: Hanabi plot Gene saturation using human embryonic stem cells (hESCs).**

The x-axis shows total counts, and y-axis shows the number of detected genes in 4 human embryo cells dataset.

**Supplementary Figure 17: Hanabi plot Gene saturation using mouse PBMC and adipocyte cells**

The x-axis shows total counts, and y-axis shows the number of detected genes in 2 mouse PBMC and adipocyte dataset.

**Supplementary Figure 18: Hanabi plot Gene saturation using human MCF10A, PBMC and HEK & 3T3 cells**

The x-axis shows total counts, and y-axis shows the number of detected genes in in different human cells dataset.

**Supplementary Figure 19: Classification of the typical and skewed coverage distribution using mouse embryonic stem cells (mESCs).**

Application of the QC methods. Left chart filter cells with low-input reads, middle chart perform trimmed clustering on the coverage matrix of the cell with high read count. A proportion of the most outlying observations is trimmed (the skewed cells). This results into two sets of cell: Typical cells with normal gene coverage and skewed cells with skewed coverage distribution. Right chart, the classification of cells was validated with the housekeeping genes (boxplot of the expression of the house keeping genes of the typical vs. skewed cells).

**Supplementary Figure 20: Classification of the typical and skewed coverage distribution using mouse CD4 T cells.**

Application of the QC methods. Left chart filter cells with low-input reads, middle chart perform trimmed clustering on the coverage matrix of the cell with high read count. A proportion of the most outlying observations is trimmed (the skewed cells). This results into two sets of cell: Typical cells with normal gene coverage and skewed cells with skewed coverage distribution. Right chart, the classification of cells was validated with the housekeeping genes (boxplot of the expression of the house keeping genes of the typical vs. skewed cells).

**Supplementary Figure 21: Classification of the typical and skewed coverage distribution using mouse fibroblast cells.**

Application of the QC methods. Left chart filter cells with low-input reads, middle chart performs trimmed clustering on the coverage matrix of the cell with high read count. A proportion of the most outlying observations is trimmed (the skewed cells). This results into two sets of cell: Typical cells with normal gene coverage and skewed cells with skewed coverage distribution. Right chart, the classification of cells was validated with the housekeeping genes (boxplot of the expression of the house keeping genes of the typical vs. skewed cells).

**Supplementary Figure 22: Classification of the typical and skewed coverage distribution using mouse hematopoietic stem cells (mHSCs)**

Application of the QC methods. Left chart filter cells with low-input reads, middle chart performs trimmed clustering on the coverage matrix of the cell with high read count. A proportion of the most outlying observations is trimmed (the skewed cells). This results into two sets of cell: Typical cells with normal gene coverage and skewed cells with skewed coverage distribution. Right panel, the classification of cells was validated with the housekeeping genes (boxplot of the expression of the house keeping genes of the typical vs. skewed cells).

**Supplementary Figure 23: Classification of the typical and skewed coverage distribution using human embryonic stem cells (hESCs).**

Application of the QC methods. Left chart filter cells with low-input reads, middle chart performs trimmed clustering on the coverage matrix of the cell with high read count. A proportion of the most outlying observations is trimmed (the skewed cells). This results into two sets of cell: Typical cells with normal gene coverage and skewed cells with skewed coverage distribution. Right chart, the classification of cells was validated with the housekeeping genes (boxplot of the expression of the house keeping genes of the typical vs. skewed cells).

**Supplementary Figure 24: Classification of the typical and skewed coverage distribution using human MCF10A and HEK & 3T3 mix cells.**

Application of the QC methods. Left chart filter cells with low-input reads, middle chart performs trimmed clustering on the coverage matrix of the cell with high read count. A proportion of the most outlying observations is trimmed (the skewed cells). This results into two sets of cell: Typical cells with normal gene coverage and skewed cells with skewed coverage distribution. Right chart, the classification of cells was validated with the housekeeping genes (boxplot of the expression of the house keeping genes of the typical vs. skewed cells).

**Supplementary Figure 25: Workflow for scRNA-seq data processing with quality assessments.**

Flow diagram summarizing the Workflow for skewness-based quality assessment of single-cell RNA-Seq experiment. Datasets are collected from three sources, in-house, public data, and 10xGenomics web resource. The figure illustrates the computation analysis tasks, and the result from each task.

**Supplementary Notes: Online resources**

