## Supplementary Figures for "Quality assessment of single-cell RNA sequencing data by coverage skewness analysis"

**Supplementary Fig. 1 | Gene coverage skewness and variation in expression among single-cell RNA-Seq protocols using mouse CD4 T cells**

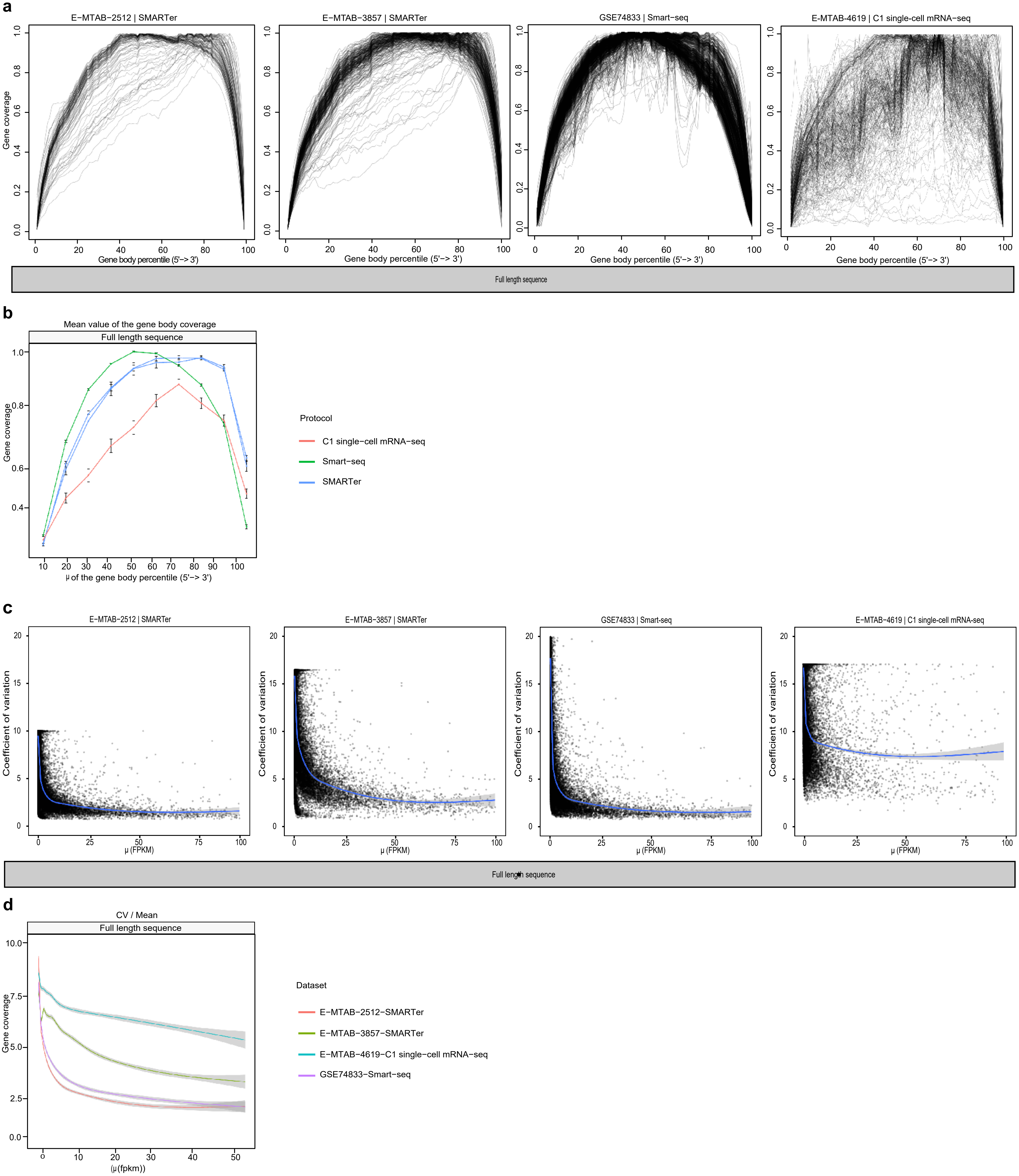

**Supplementary Fig. 2 | Gene coverage skewness and variation in expression among single-cell RNA-Seq protocols using mouse fibroblast cells**

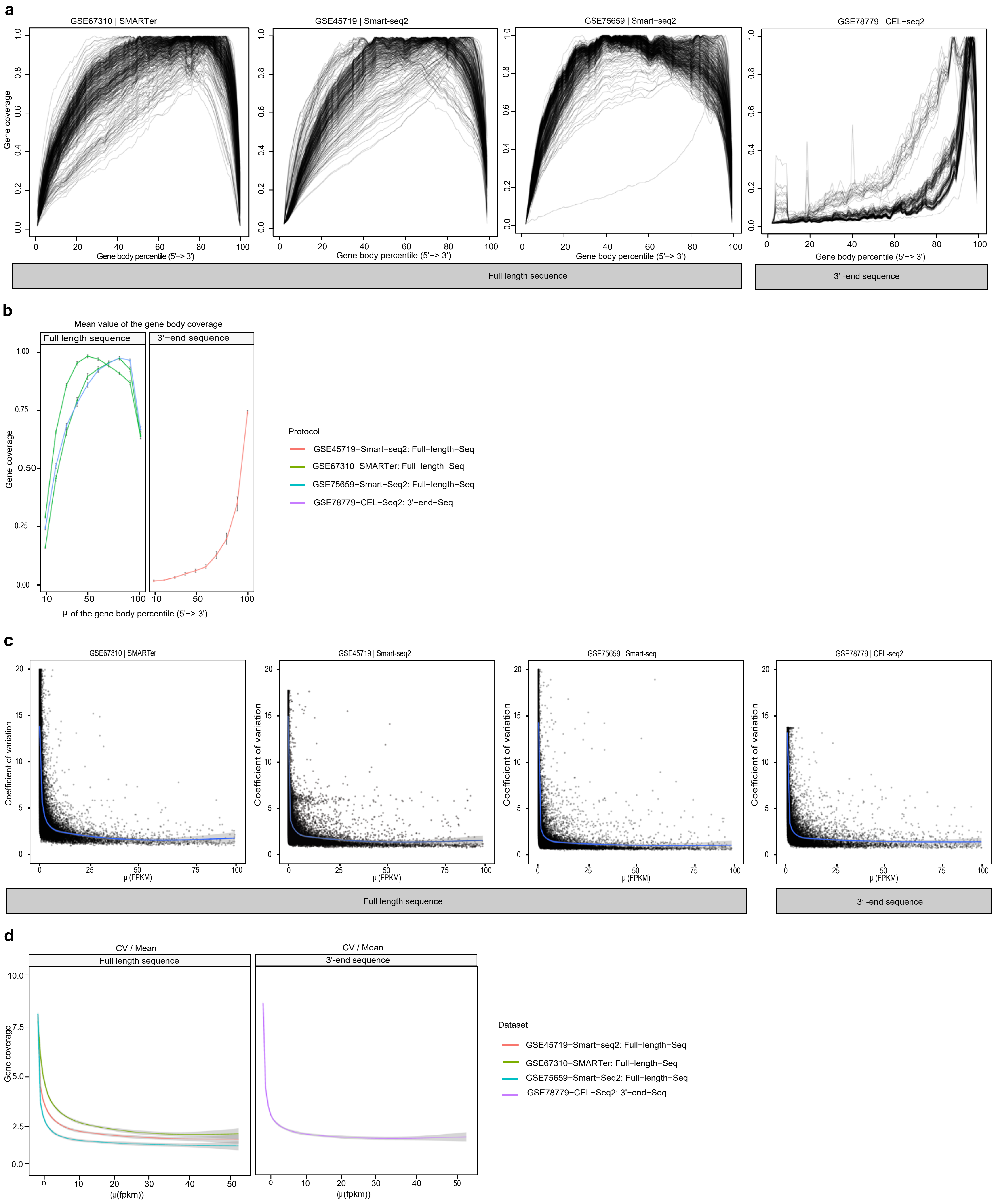

**Supplementary Fig. 3 | Gene coverage skewness and variation in expression among single-cell RNA-Seq protocols using mouse hematopoietic stem cells (HSCs)**

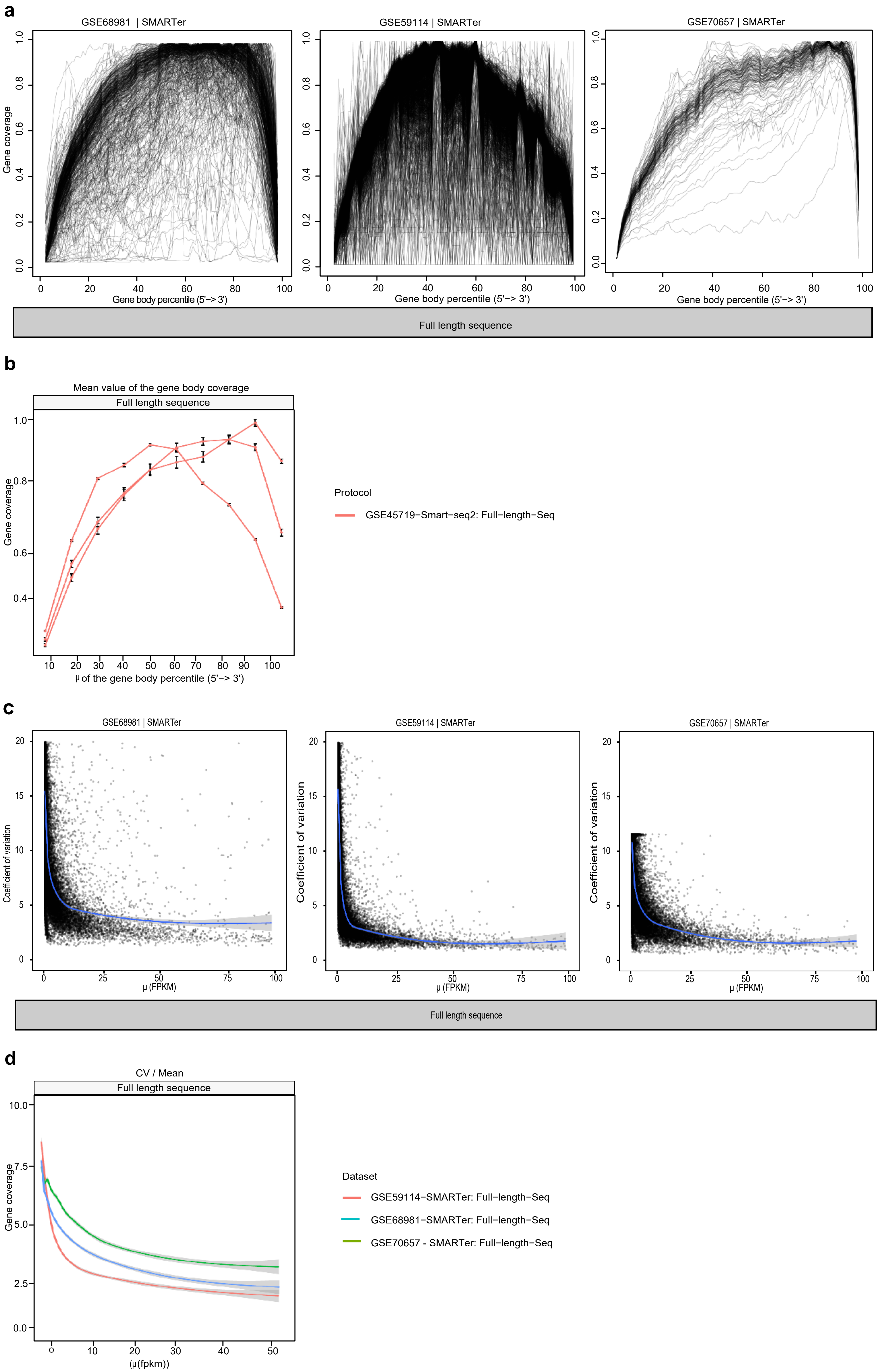

**Supplementary Fig. 4 | Gene coverage skewness and variation in expression among single-cell RNA-Seq protocols using human embryonic stem cells (hESCs)**

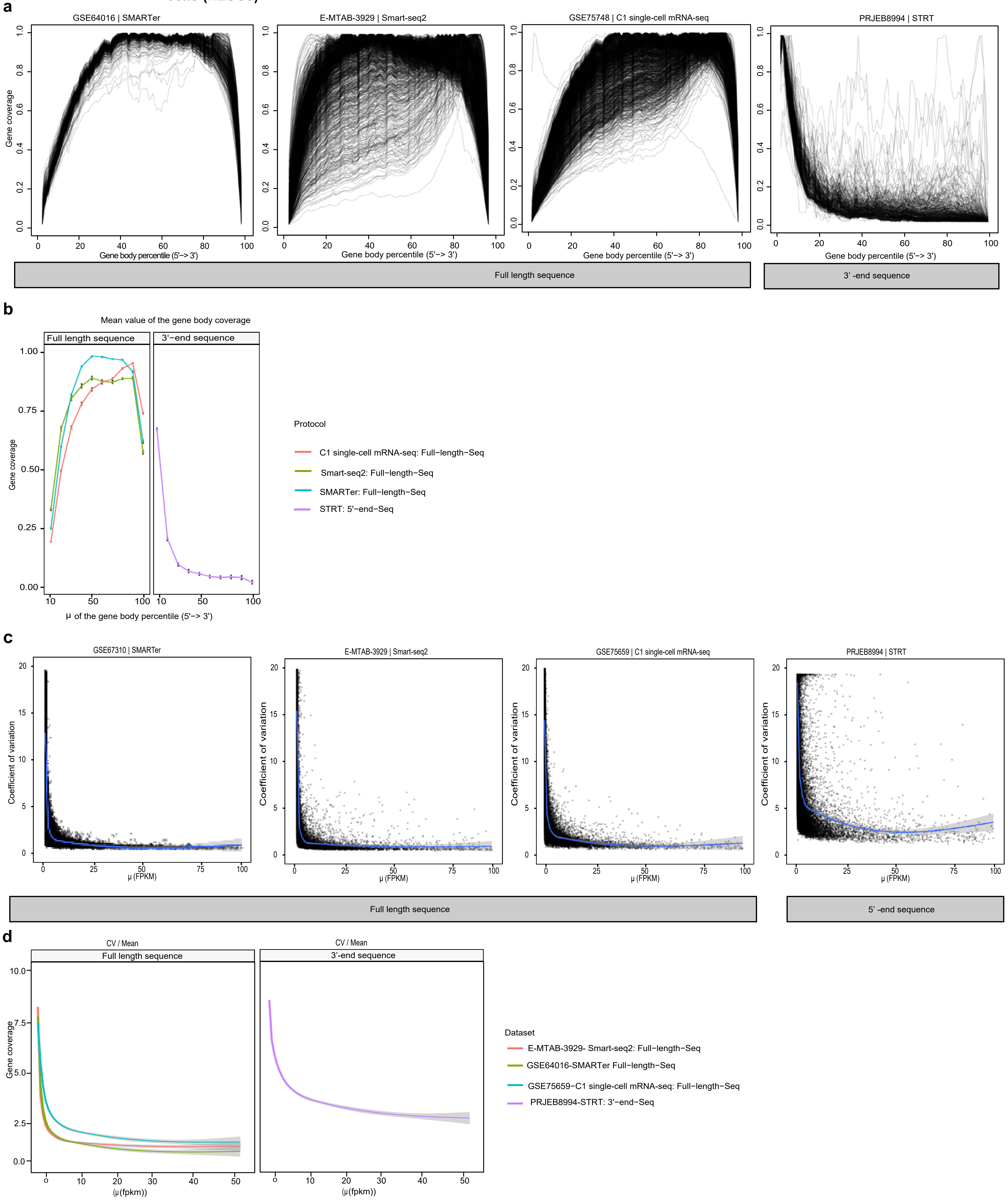

**Supplementary Fig. 5 | Gene coverage skewness and variation in expression among single-cell RNA-Seq protocols using mouse un-matched cells**

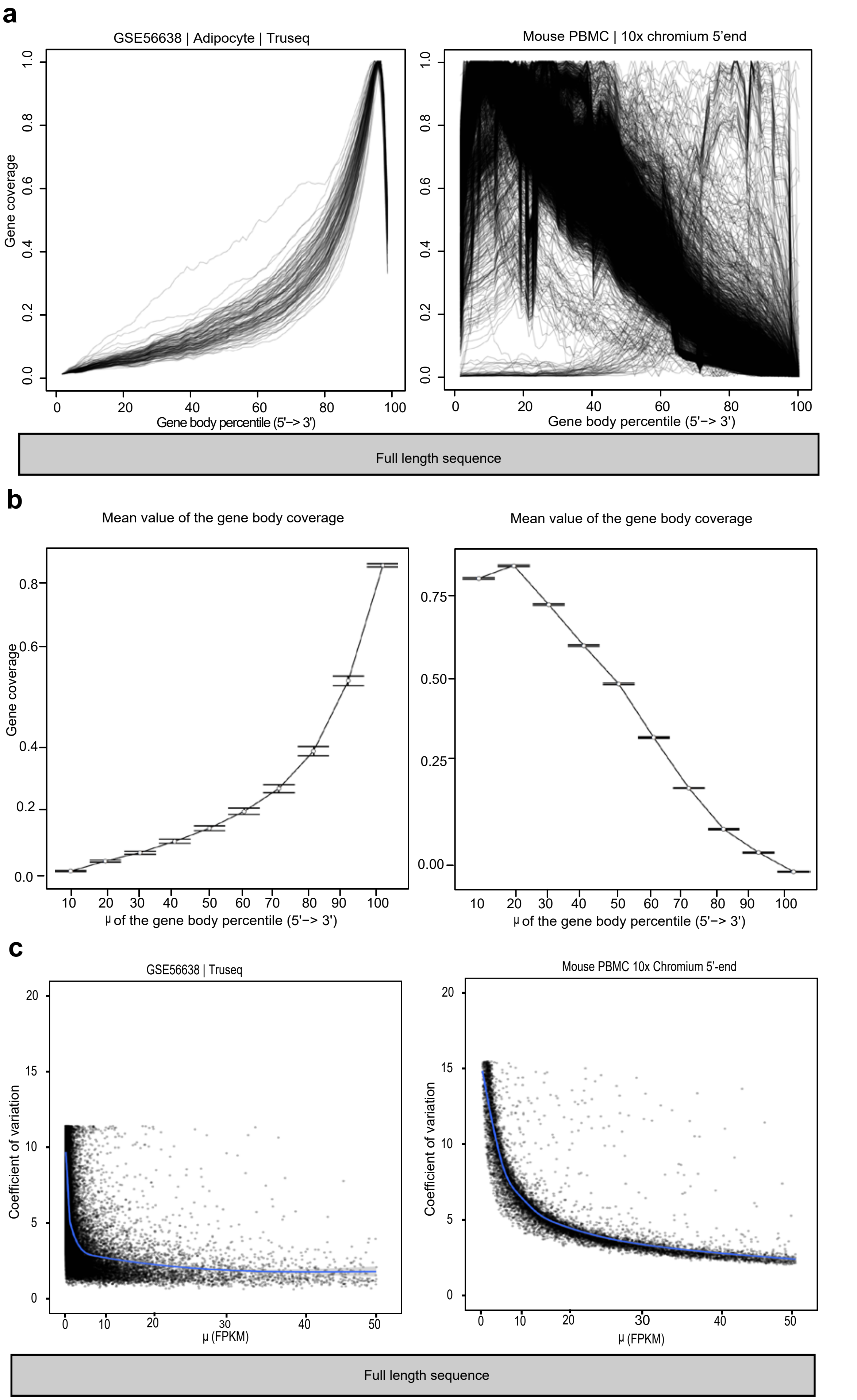

**Supplementary Fig. 6 | Gene coverage skewness and variation in expression among single-cell RNA-Seq protocols using human un-matched cells**

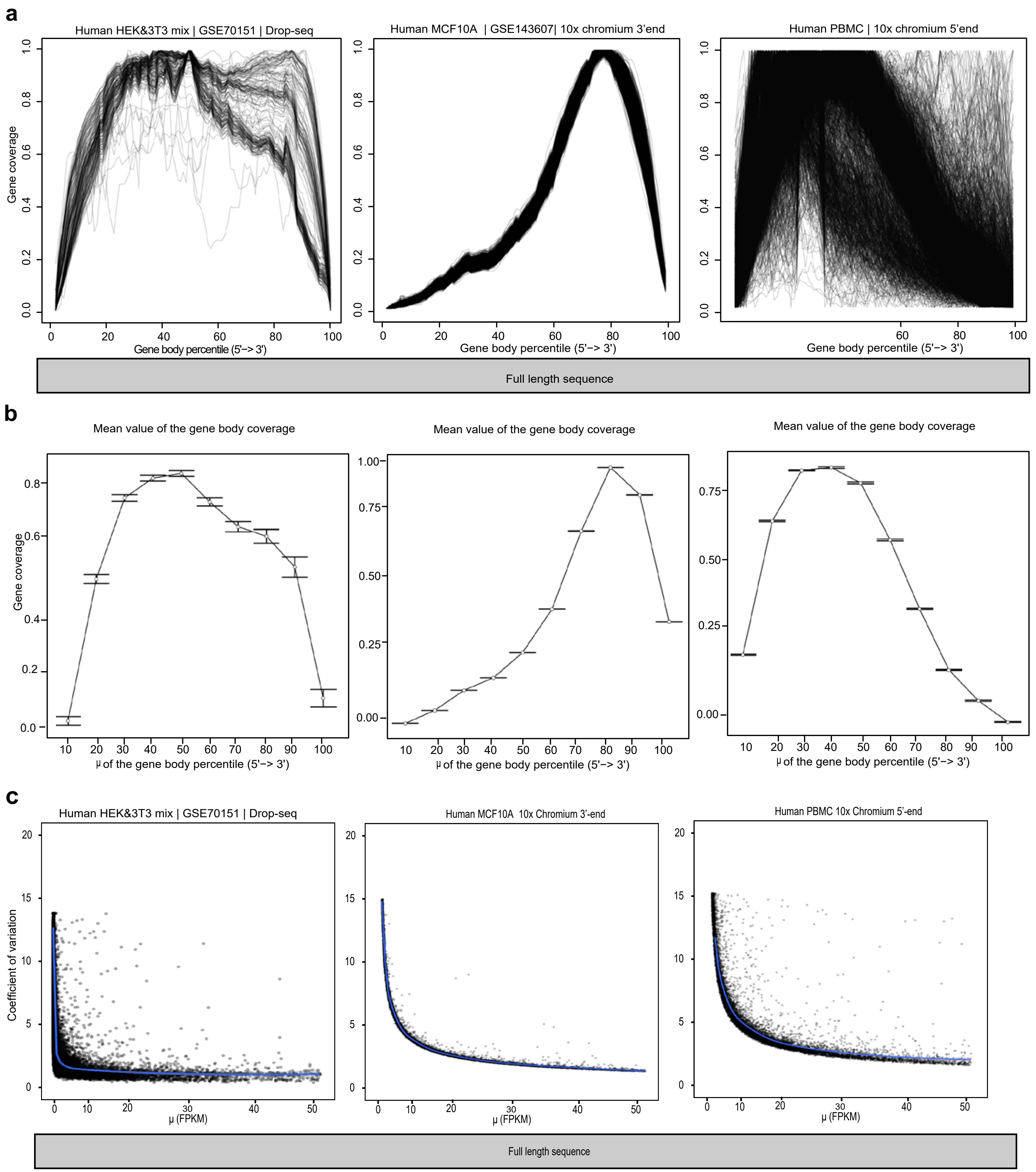

**Supplementary Fig. 7 | Dataset-to-dataset similarity of the mean expression of mouse embryonic stem cells (mESCs)**

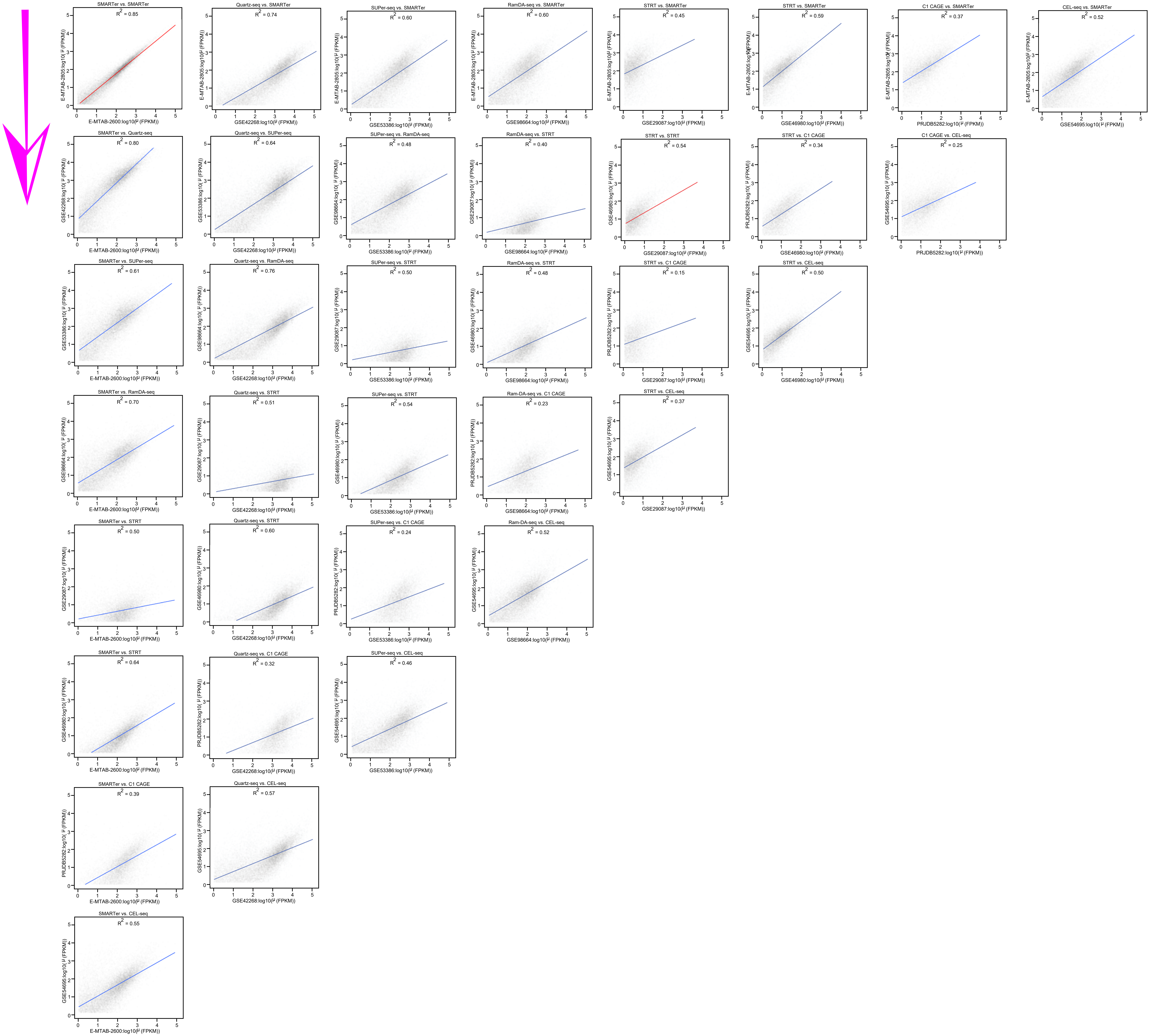

**Supplementary Fig. 8 | Dataset-to-dataset similarity of the mean expression of mouse CD4 T cells**

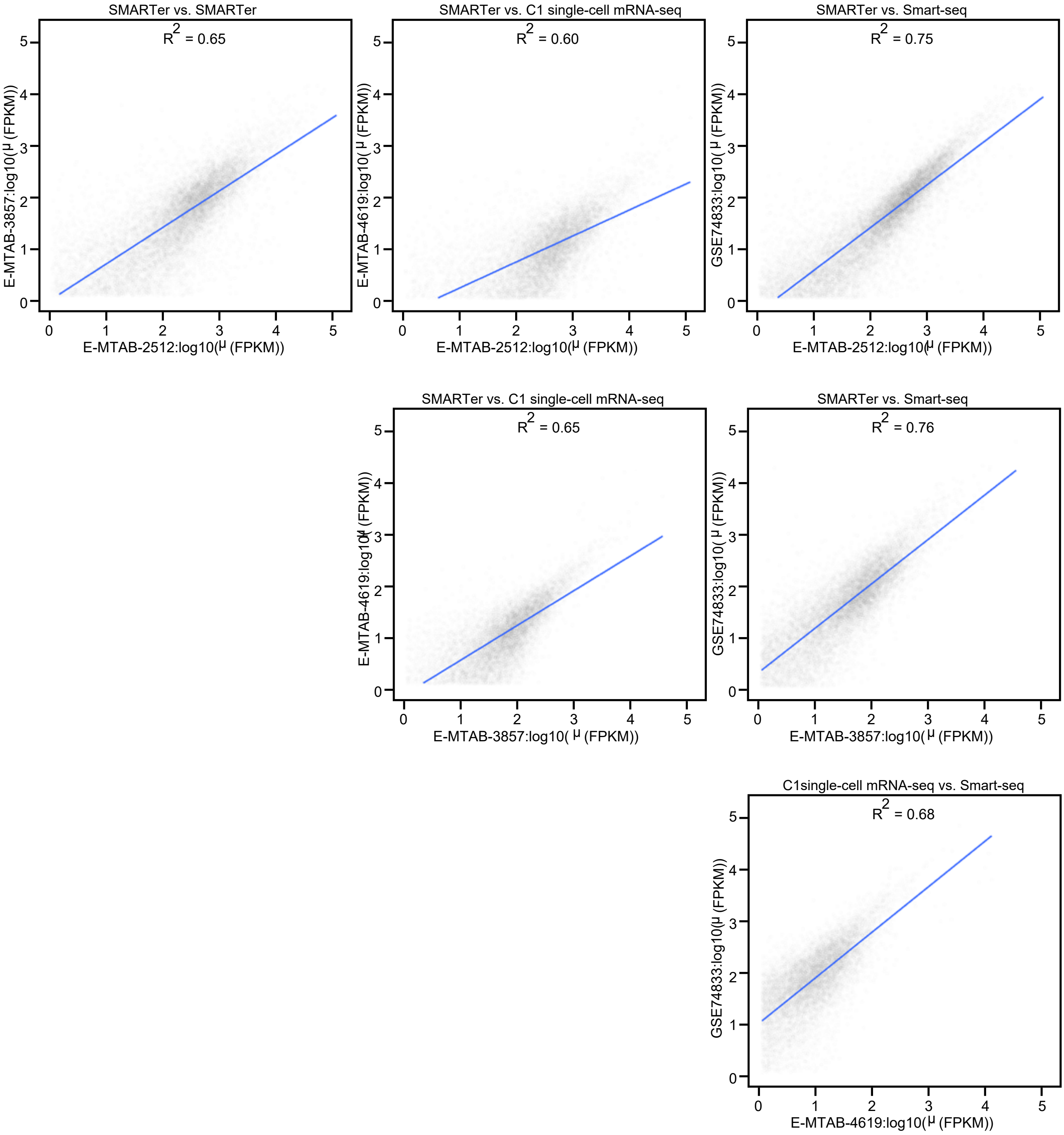

**Supplementary Fig. 9 | Dataset-to-dataset similarity of the mean expression of mouse fibroblast cells**

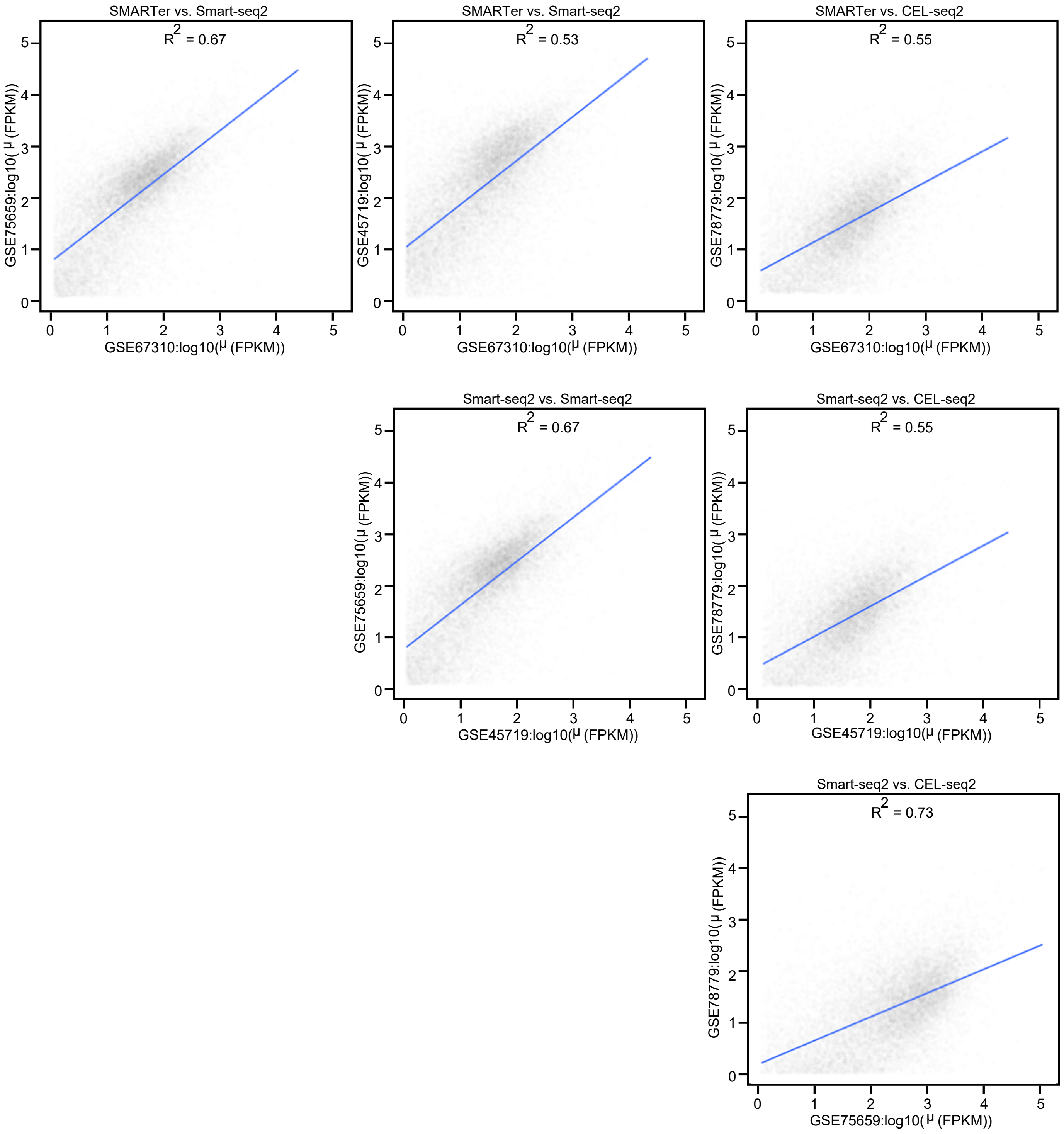

**Supplementary Fig. 10 | Dataset-to-dataset similarity of the mean expression of mouse hematopoietic stem cells (HSCs)**

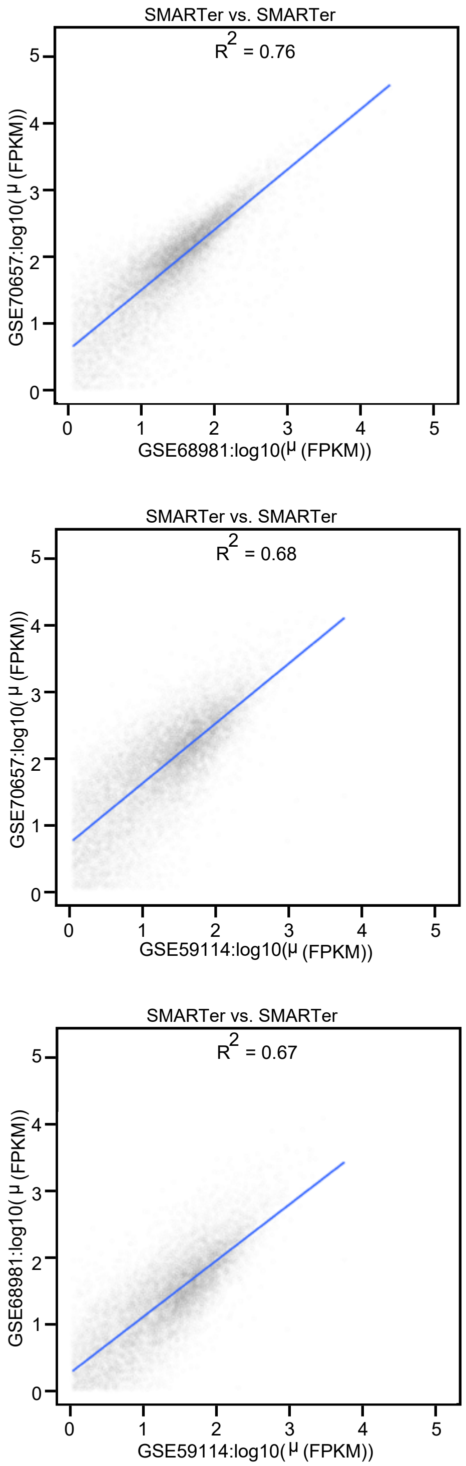

**Supplementary Fig. 11 | Dataset-to-dataset similarity of the mean expression of Human embryonic stem cells (hESCs)**

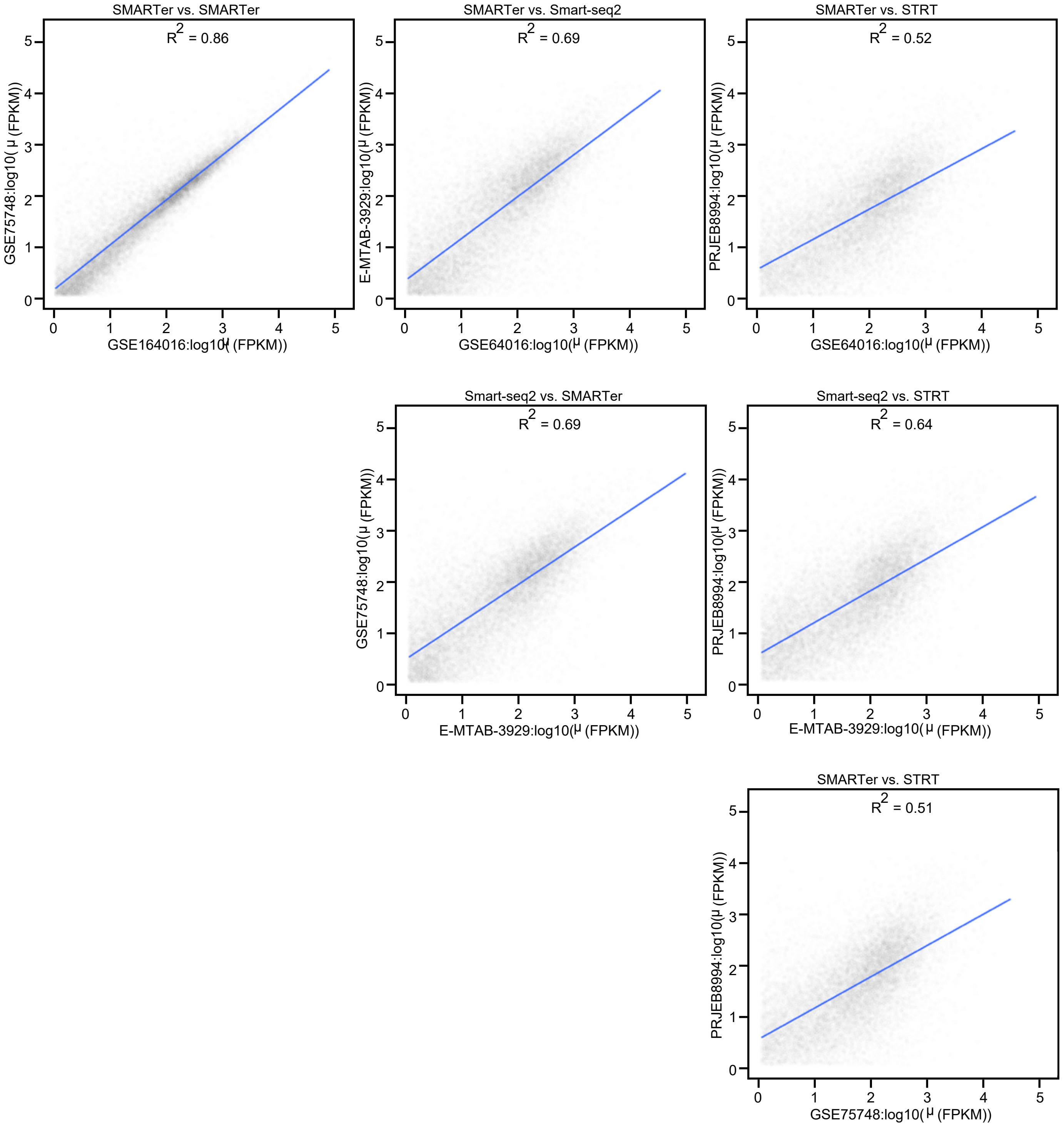

### Supplementary Fig. 12 | Hanabi plot gene saturation using mouse embryonic stem cells (mESC)

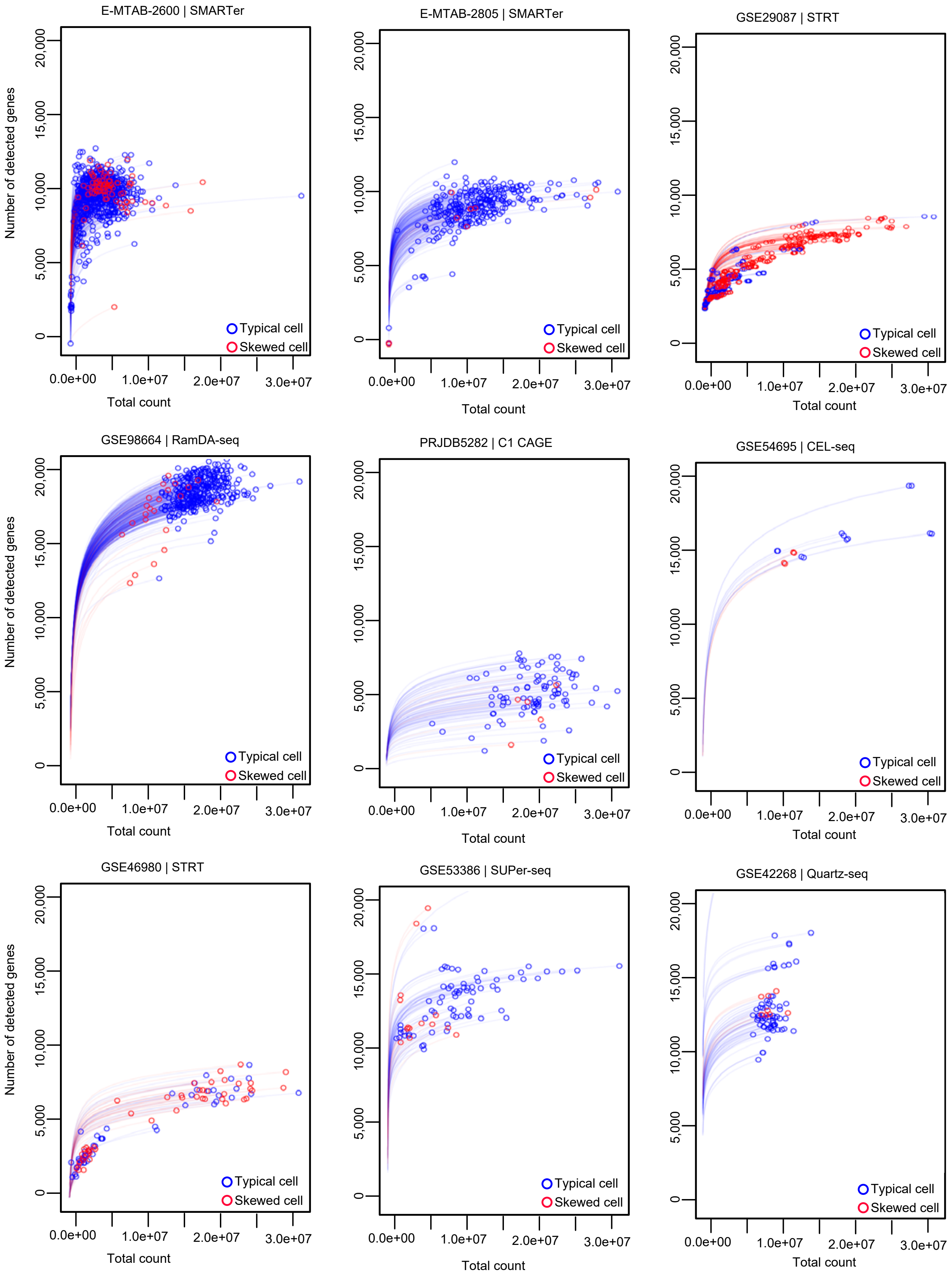

**Supplementary Fig. 13 | Hanabi plot gene saturation using mouse CD4 T cells**

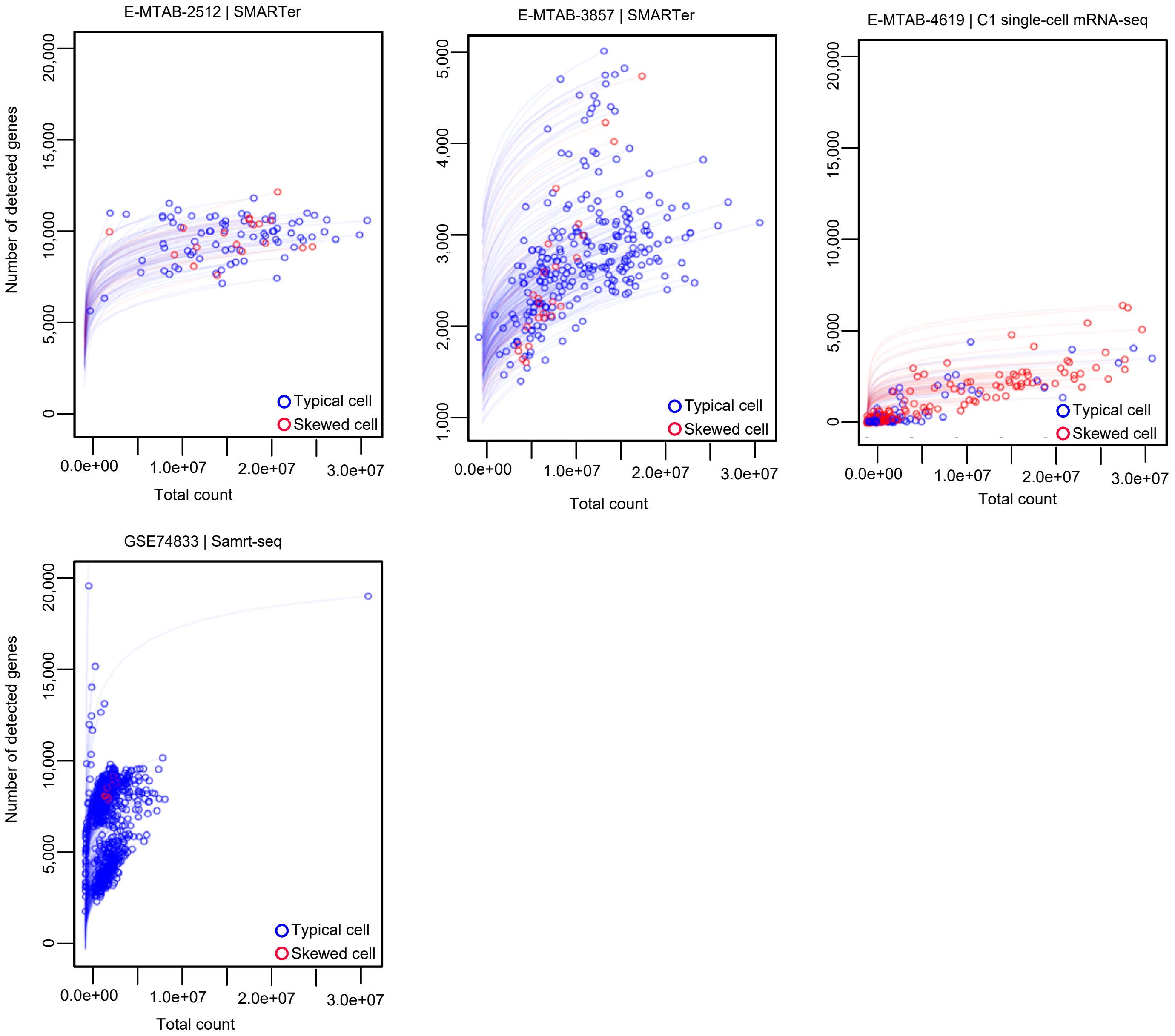

#### Supplementary Fig. 14 | Hanabi plot gene saturation using mouse fibroblast cells

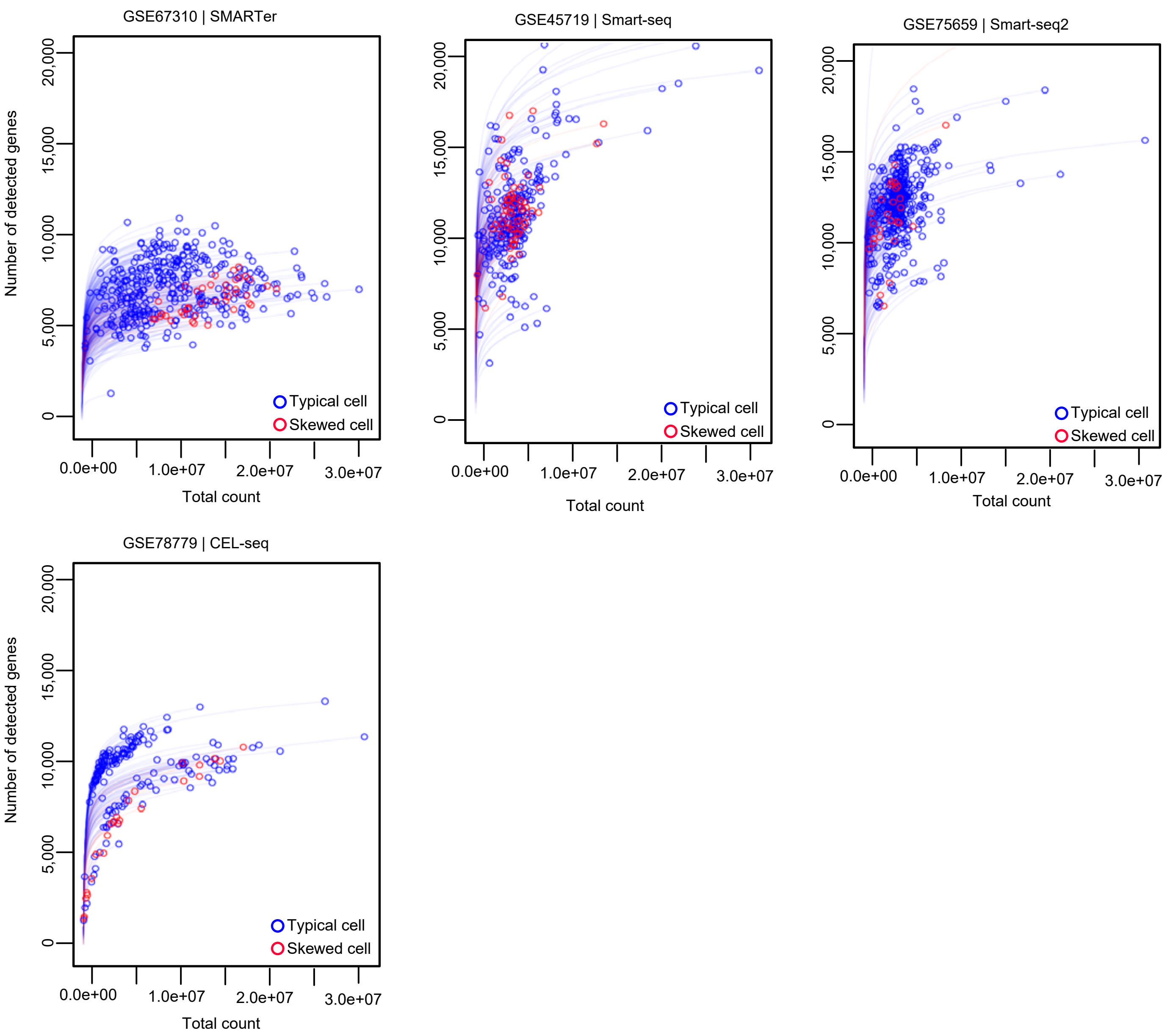

### Supplementary Fig. 15 | Hanabi plot gene saturation using mouse hematopoietic stem cells (HSCs)

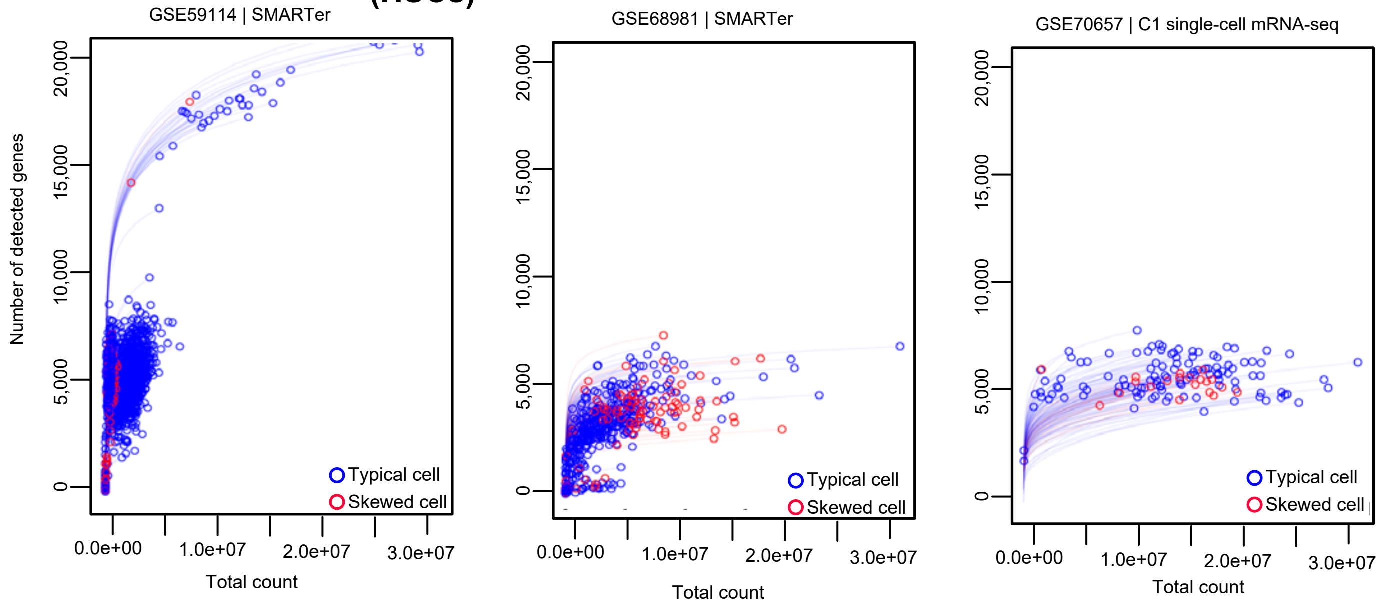

**Supplementary Fig. 16 | Hanabi plot gene saturation using human embryonic stem cells (hESCs)**

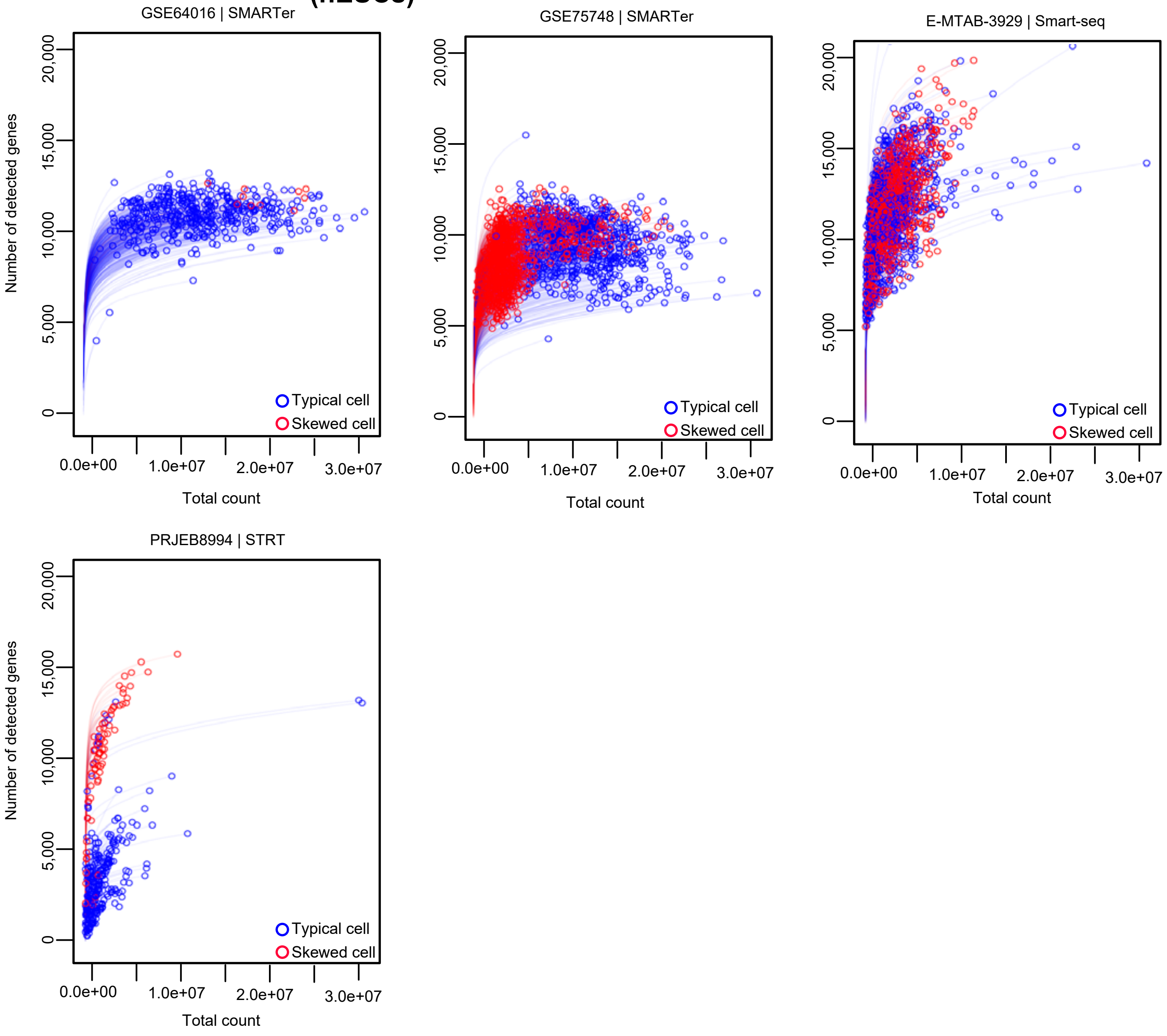

### Supplementary Fig. 17 | Hanabi plot gene saturation using mouse PBMC and adipocyte cells

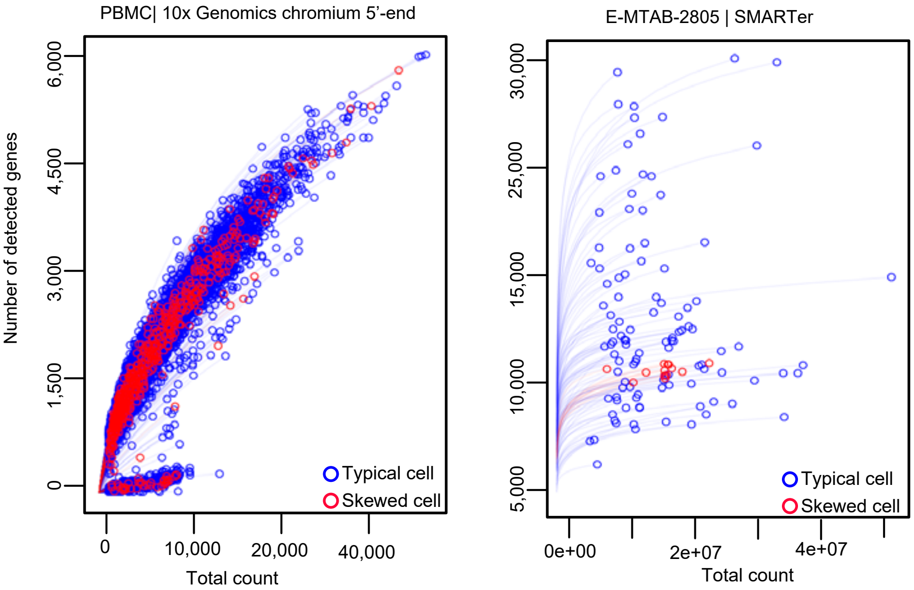

#### Supplementary Fig. 18 | Hanabi plot gene saturation using human MCF10A, PBMC and HEK & 3T3 cells

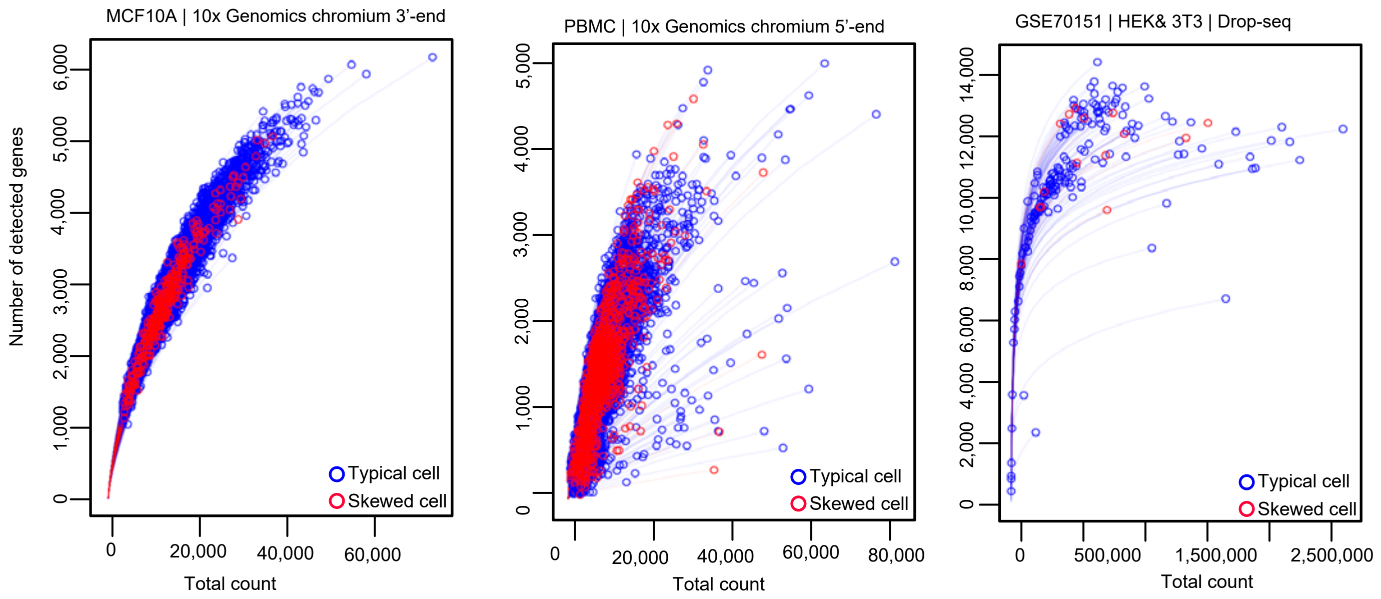

**Supplementary fig. 19 | Classification of typical and skewed coverage distribution cells using mESC**

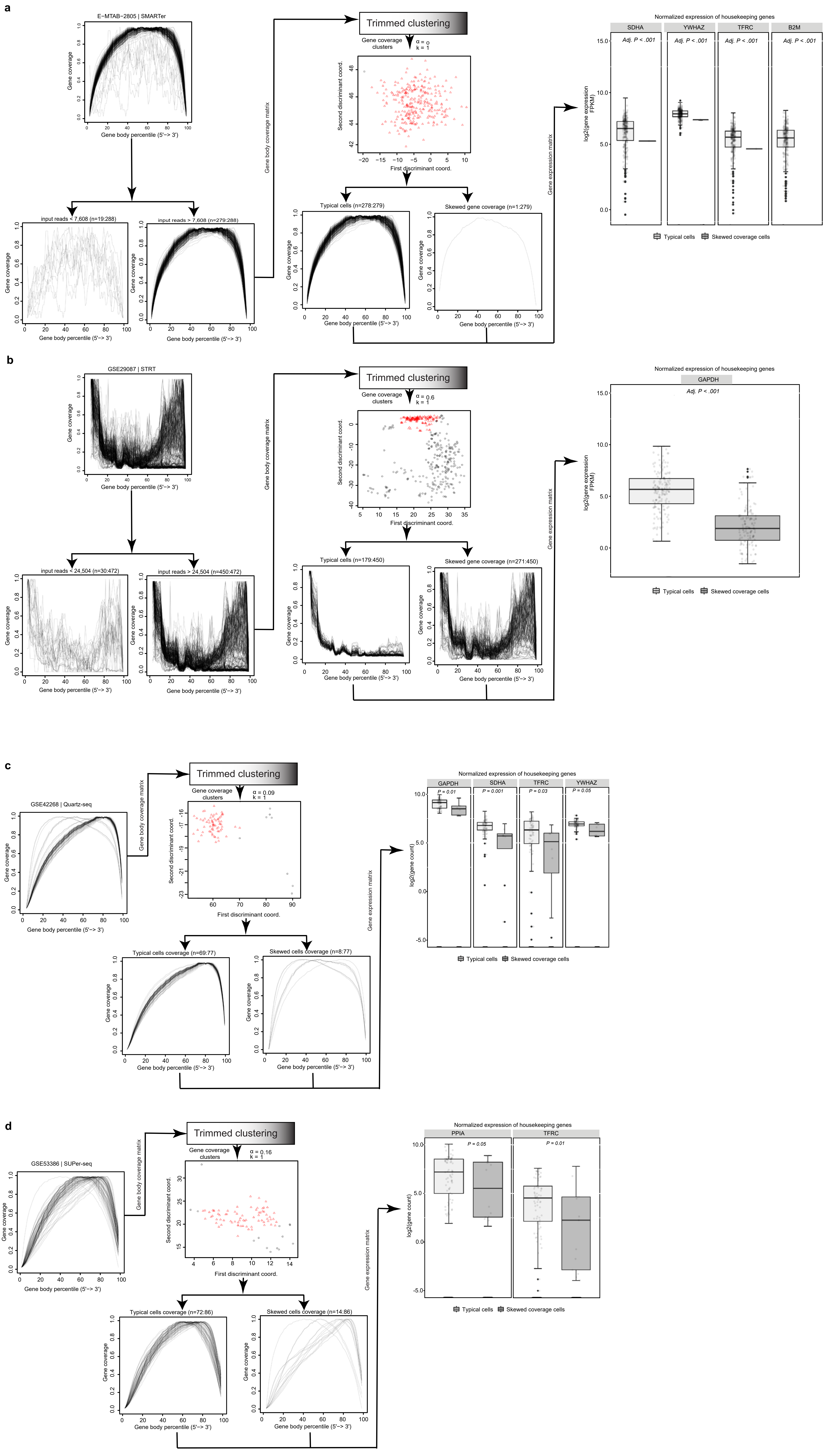

**Supplementary fig. 19 | Classification of typical and skewed coverage distribution cells using mESC**

**e**

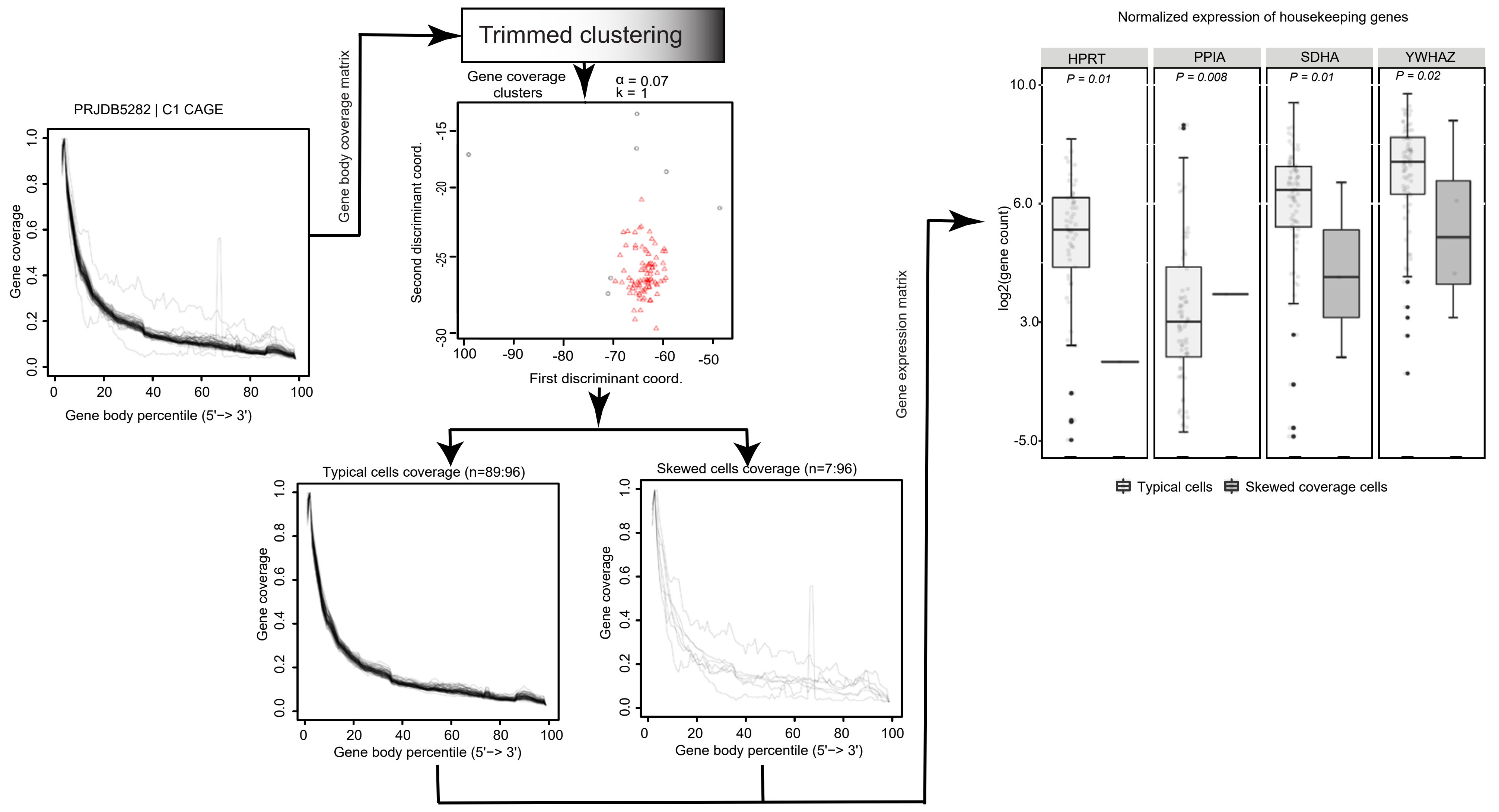


**f**

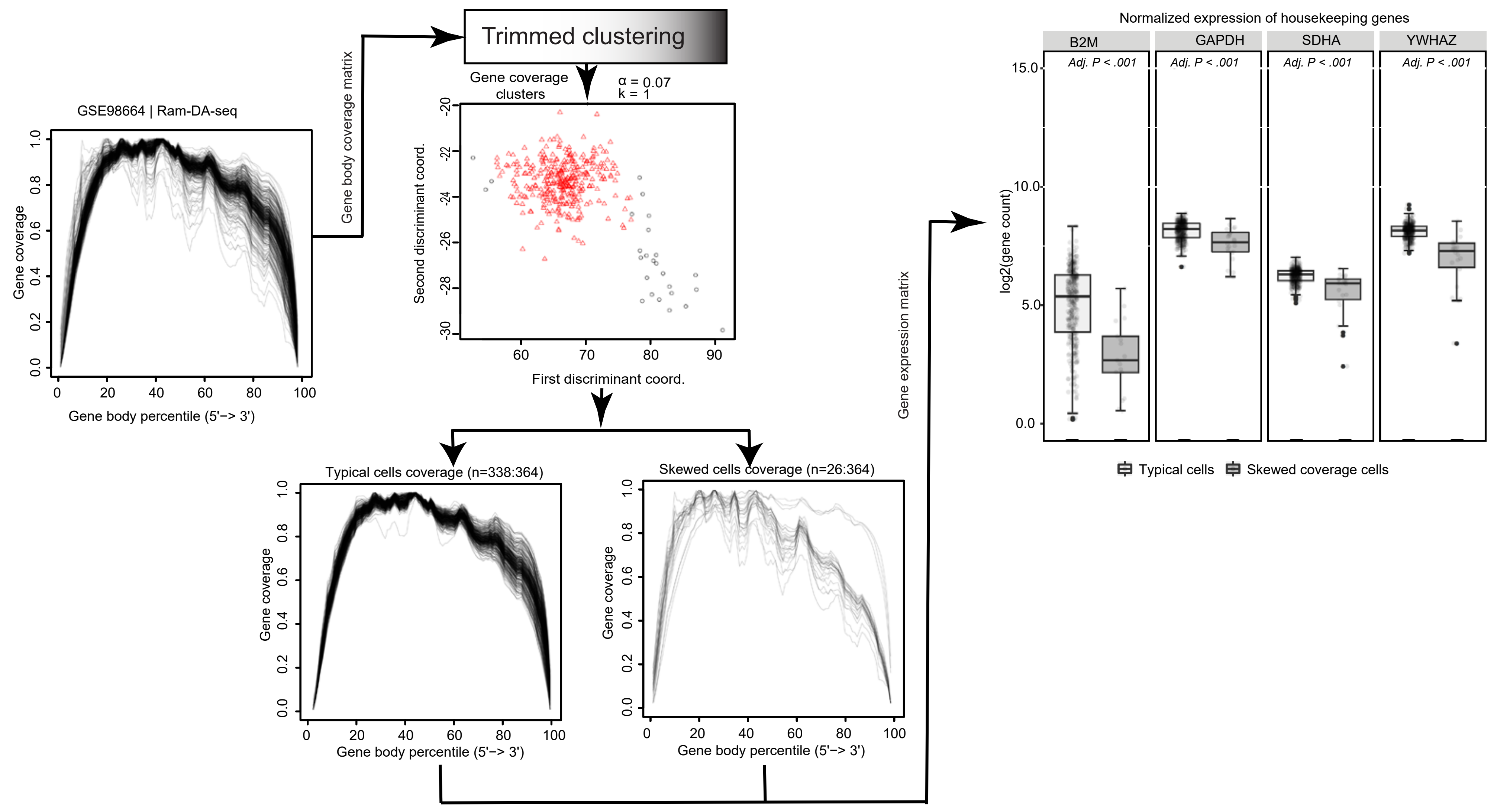


**g**

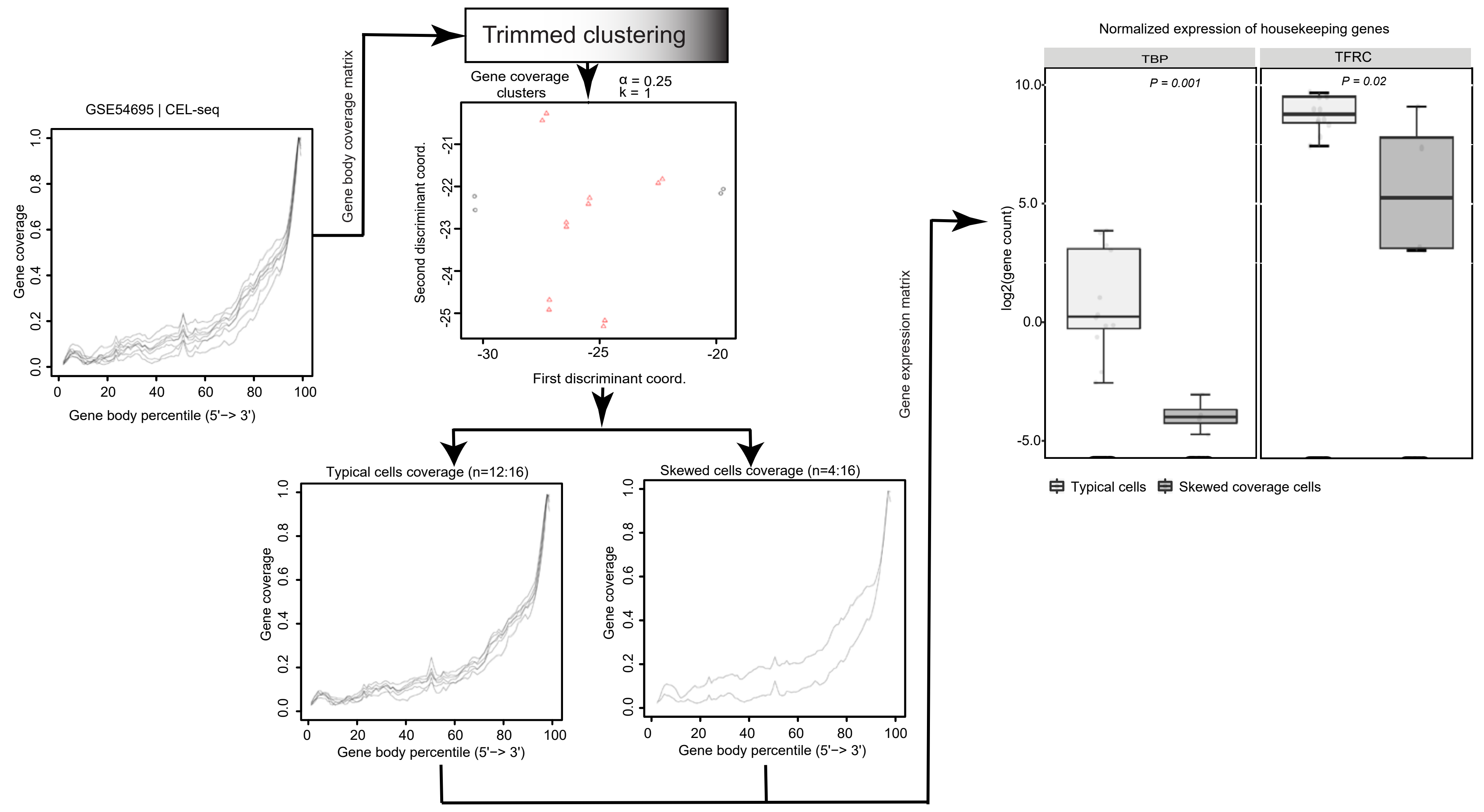

**Supplementary fig. 20 | Classification of typical and skewed coverage distribution cells using mouse CD4 T cells**

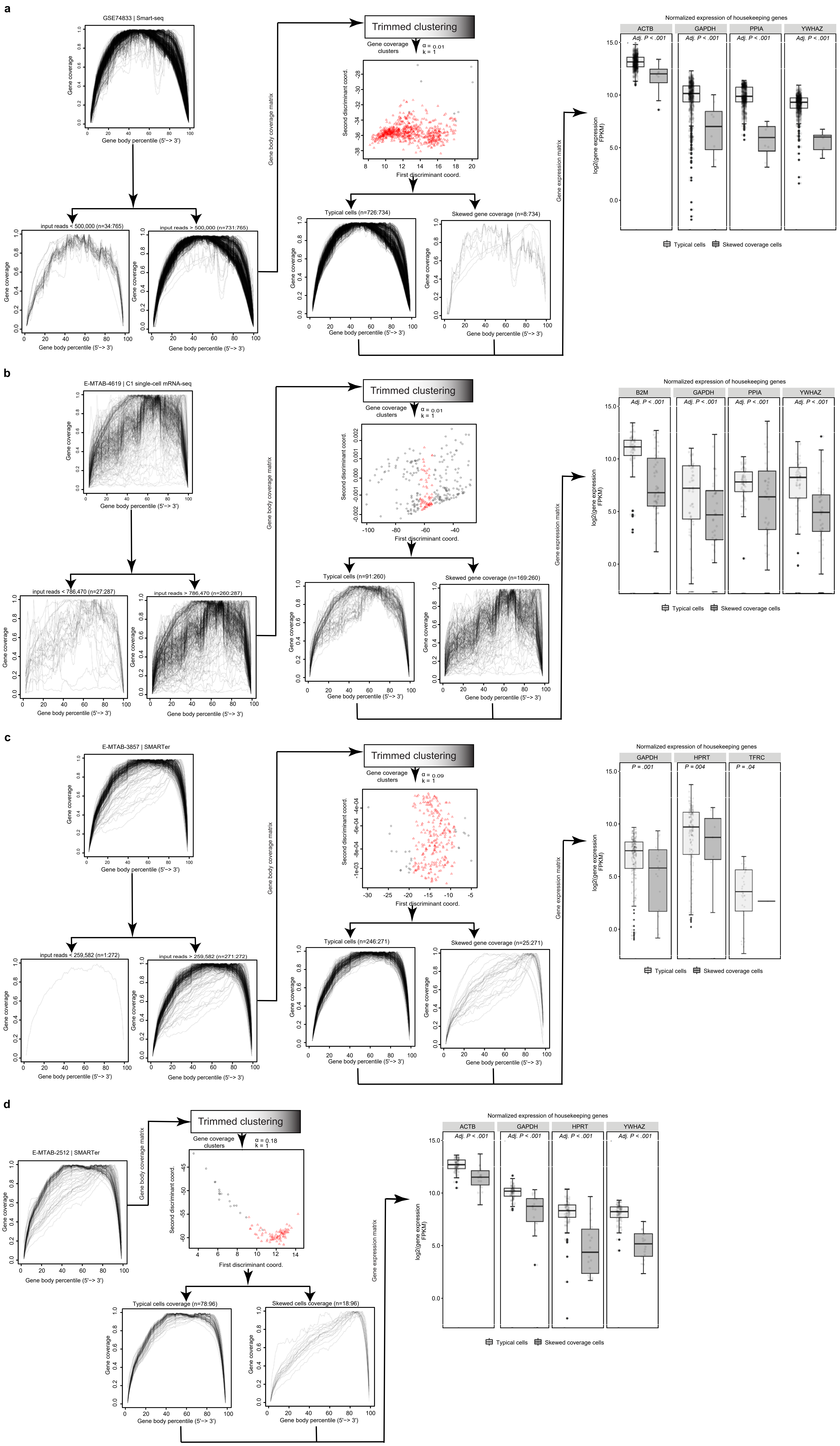

**Supplementary fig. 21 | Classification of typical and skewed coverage distribution cells using mouse fibroblast cells**

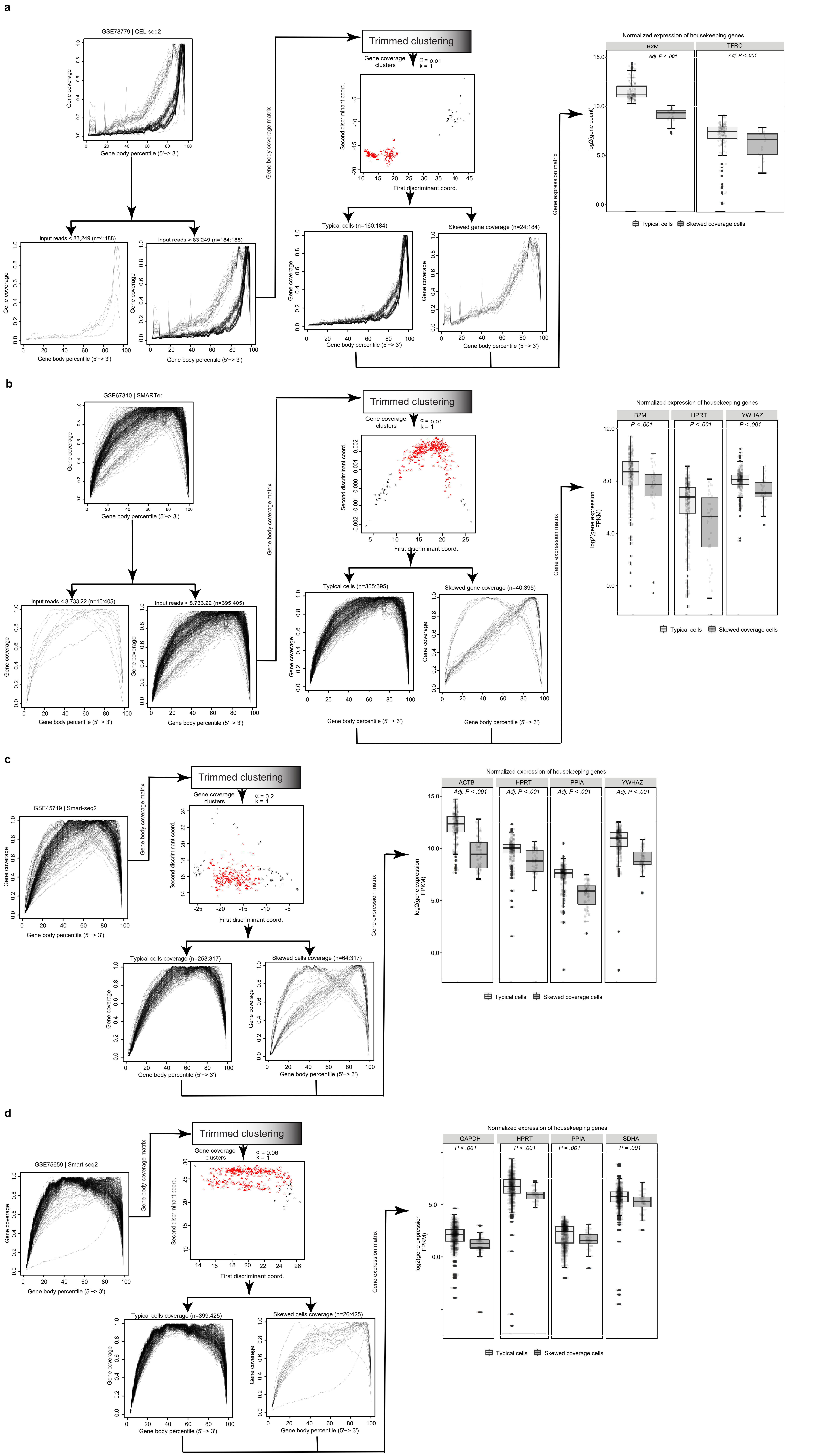

**Supplementary fig. 22 | Classification of typical and skewed coverage distribution cells using mouse hematopoietic stem cells (HSCs)**

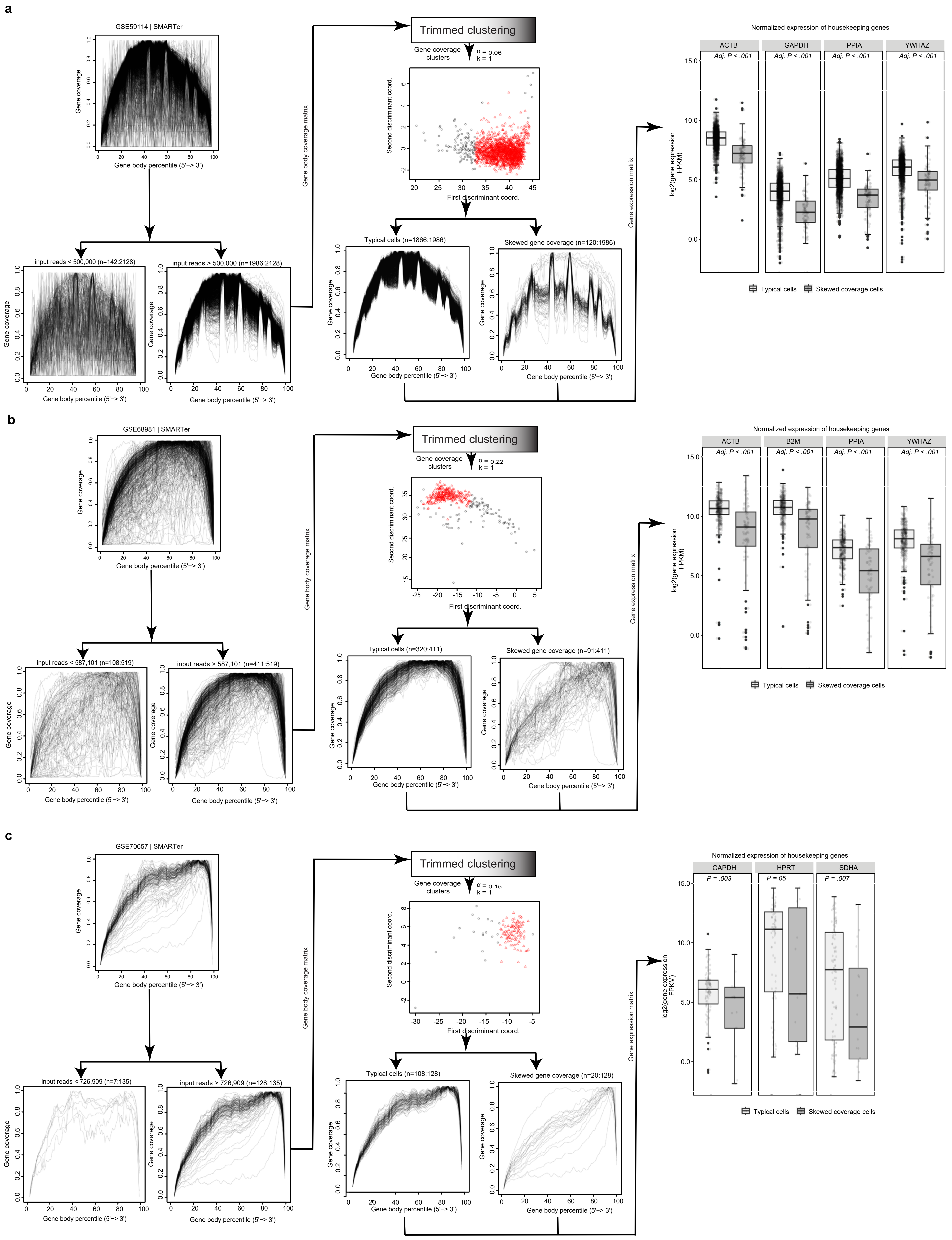

**Supplementary fig. 23 | Classification of typical and skewed coverage distribution cells using human embryonic stem cells (hESCs)**

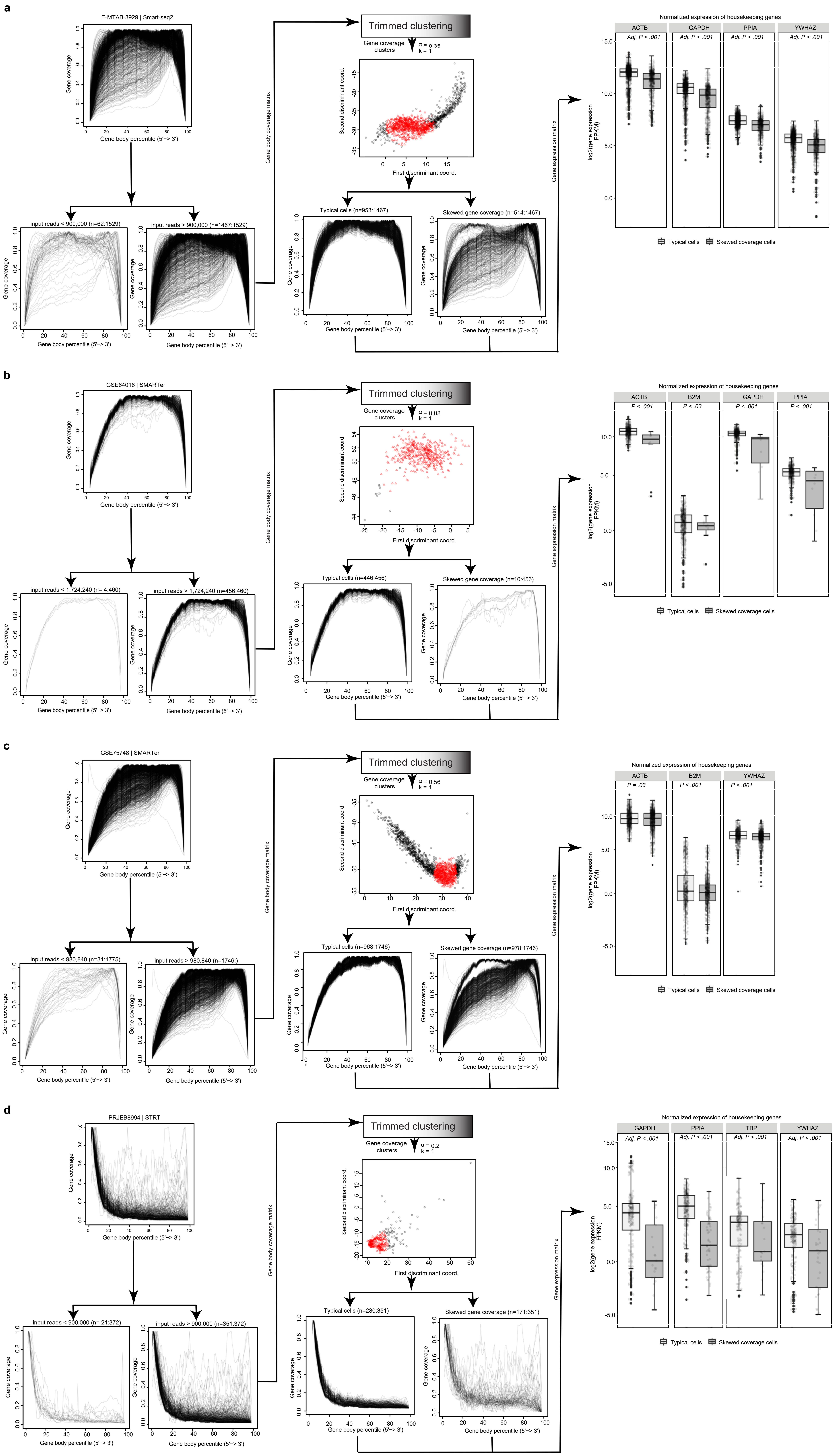

**Supplementary fig. 24 | Classification of typical and skewed coverage distribution cells using MCF10A and HEk & 3T3 mix cells**

**a**

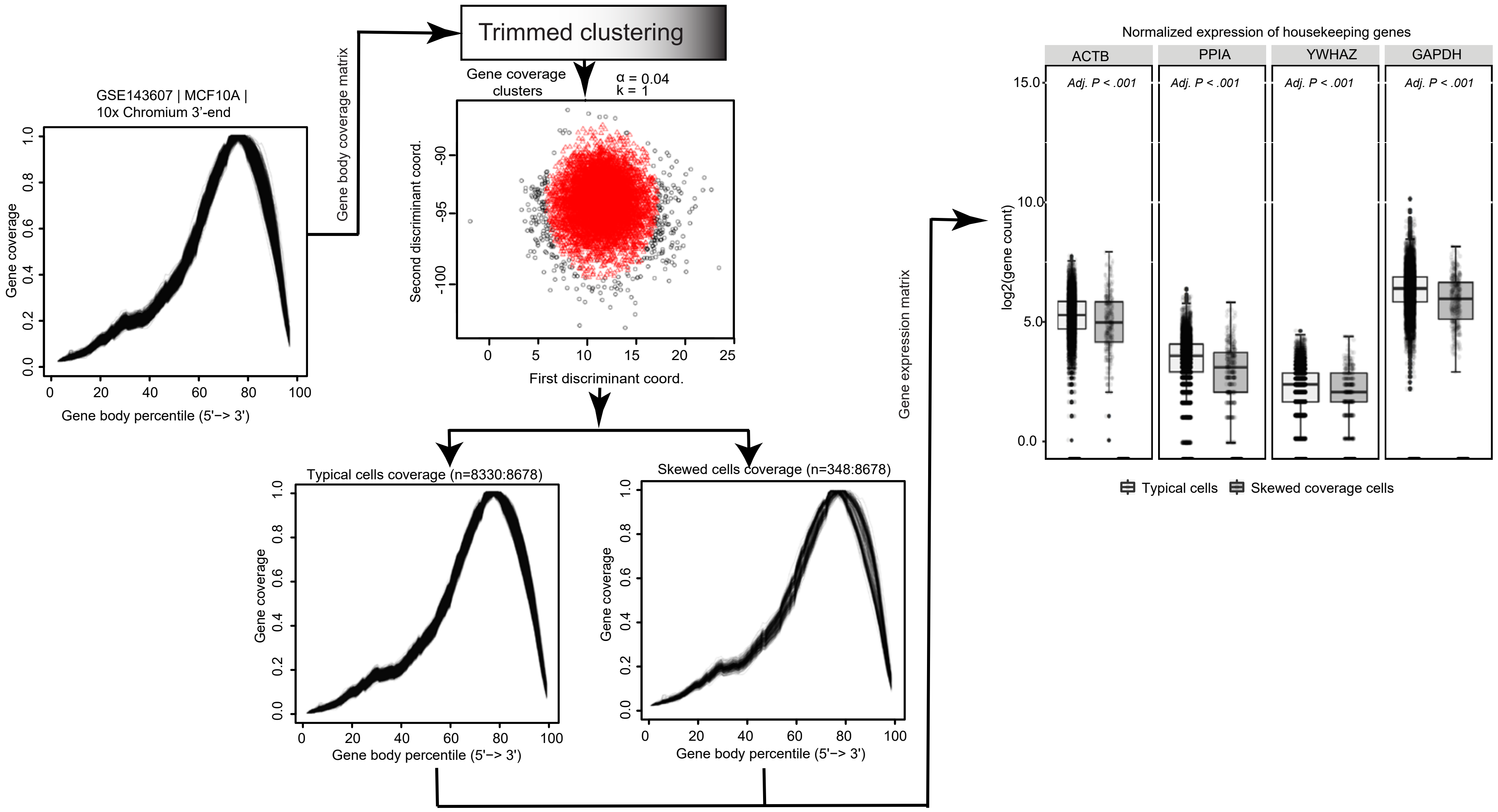


**b**

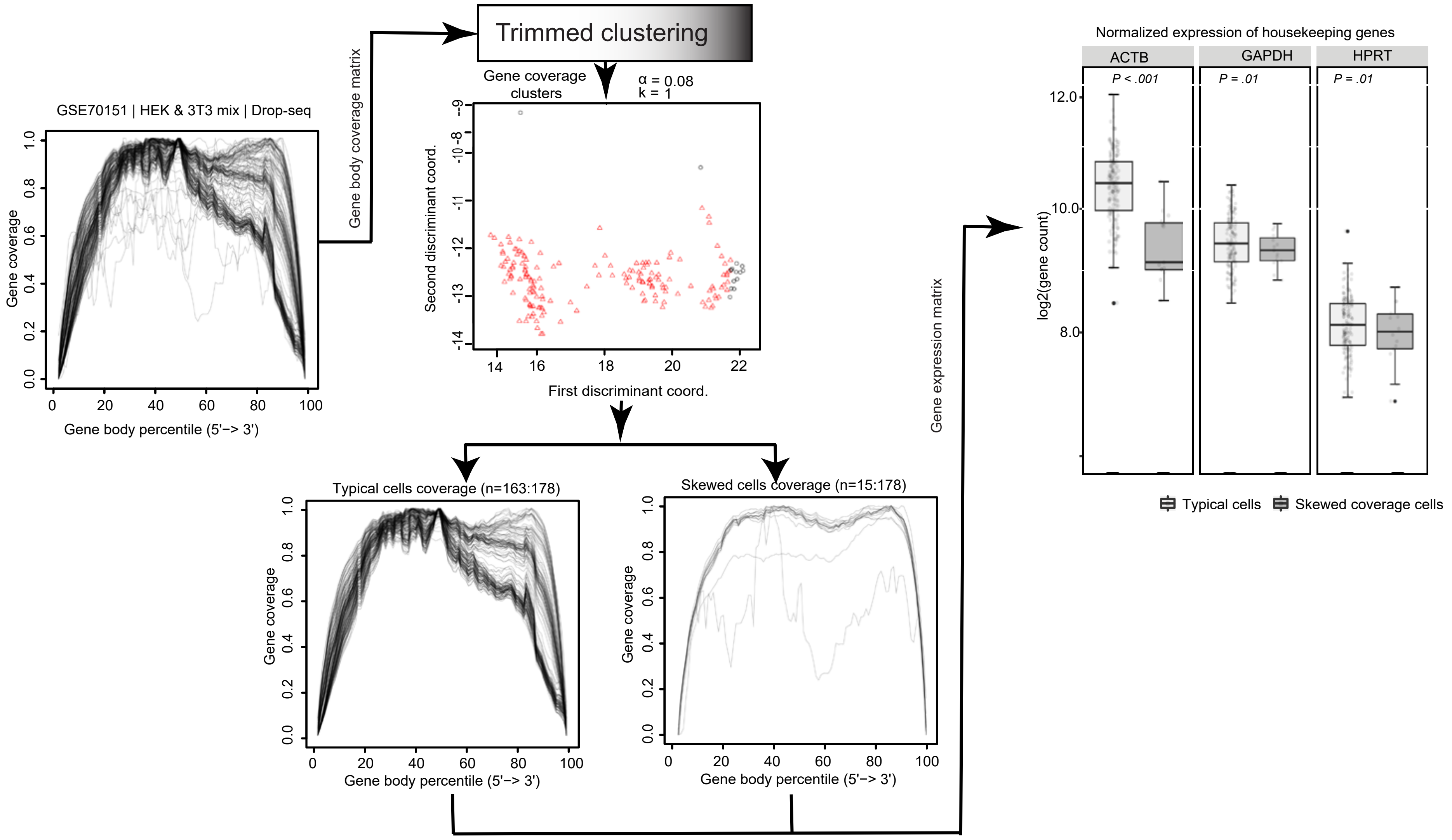

### Supplementary Fig. 25 | Workflow for scRNA-seq data processing with quality assessments

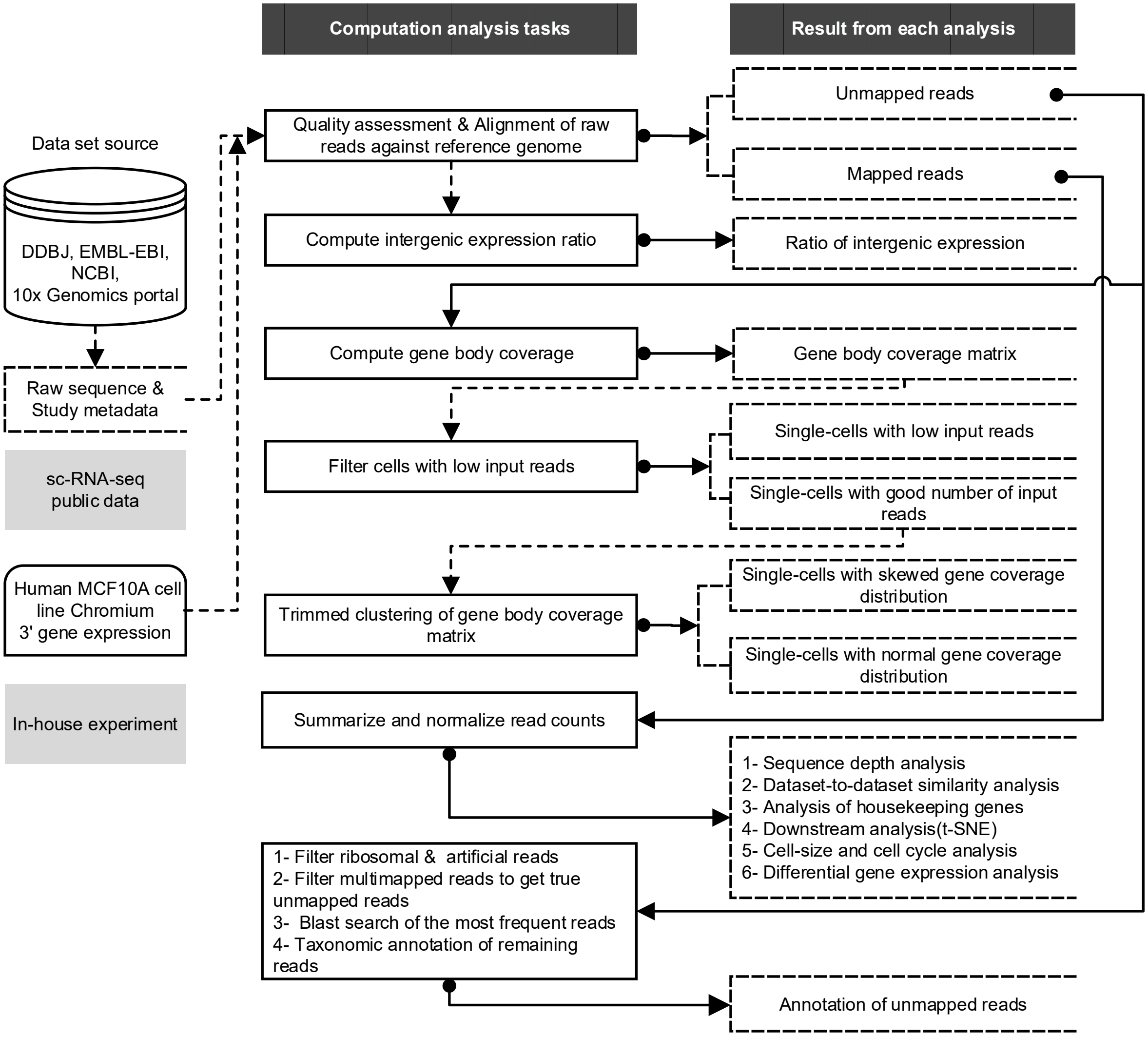
