## Supplementary Notes for "Quality assessment of single-cell RNA sequencing data by coverage skewness analysis"

TABLE OF C ONTENTS

### Software scripts and tools

1 | *Workflow for scRNA-seq data collection, primary analysis.*
The workflow and scripts used for collecting and processing scRNA-seq datasets for this manuscript was available in GitHub under https://github.com/LSBDT/SCPortalen and licensed under the Creative Commons Zero v1.0 Universal. Usage instructions and prerequisite tools are described in the README.md.

2 | *R scripts for performing SkewC method for classification of typical and skewed cells*
SkewC was implemented in R, the R markdown to use the method available with online instructions under https://github.com/LSBDT/SkewC . As described in the manuscript, the input for SkewC was the indexed BAM file and the gene model in BED format.

3 | *SkewC10X Genomics: Processing pipeline for 10XGenomics scRNA-seq dataset*
10X genomics generate pooled BAM file per each dataset, using cell barcodes to identify each cell. SkewC10X provide Docker image under https://hub.docker.com/repository/docker/moirai2/skewc10x SkewC10X split the pooled BAM file into smaller BAM files (one per each single-cell), then compute the gene body coverage. Please follow the instructions to download, setup and run the Docker.

### The interactive web-based scRNA-seq database: SCPortalen

Raw and processed scRNA-seq dataset used in this manuscript are available through online interactive web database (SCPortalen). Additionally, analysis results are provided for query, download and exploration. SCPortalen accessible at http://single-cell.clst.riken.jp/

1 | *SCPortalen overview*

The online database landing page provide access to scRNA-seq data and single-cell images. Here we'll focus on scRNA-seq data only  as shown below.

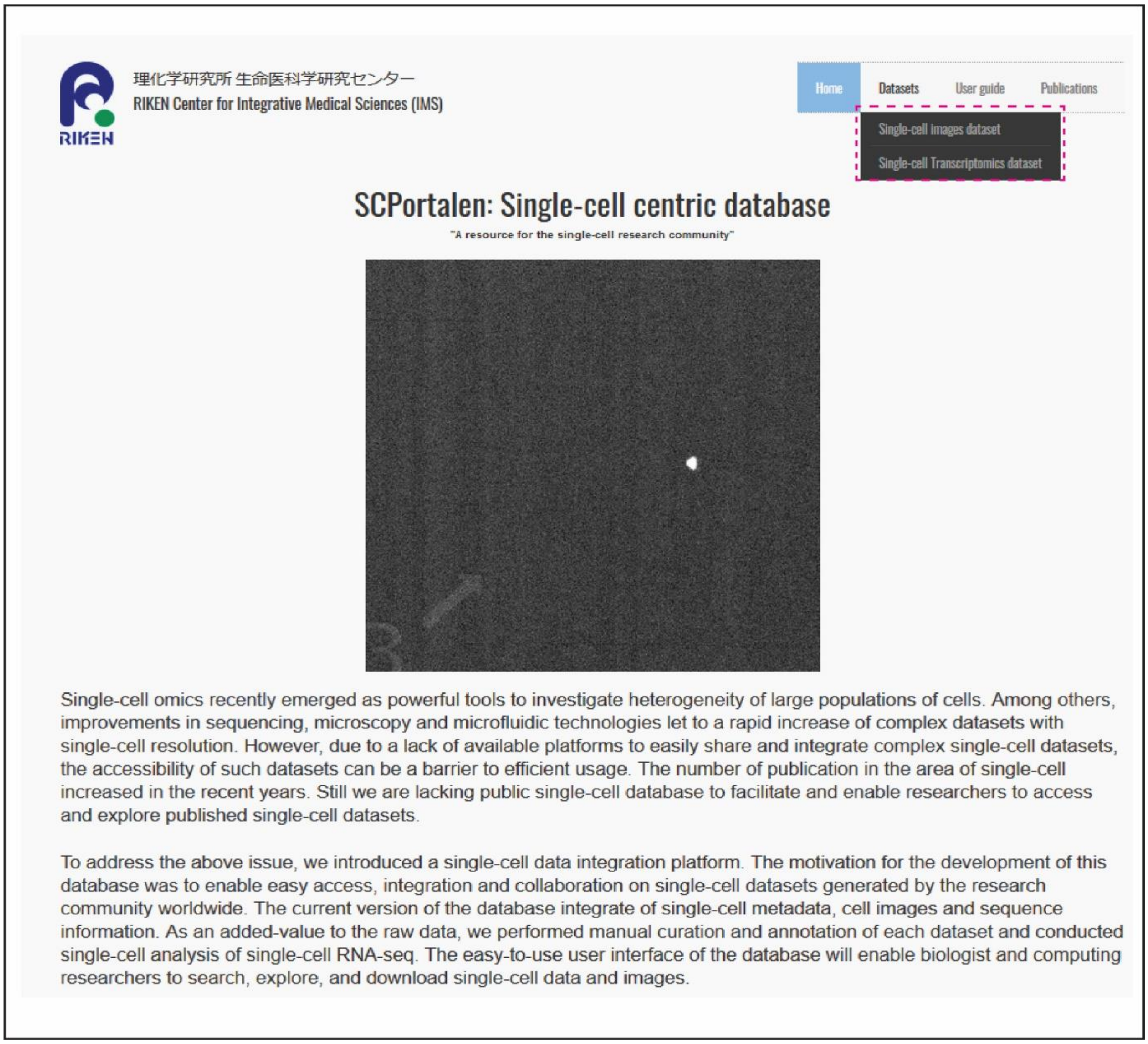


Single-cell database landing page provides access to the single-cell transcriptomics dataset and analysis.

2 | *Dataset overview*

For each scRNA-seq dataset SCPortalen provides dataset view which shows:

1. Different attributes (metadata), and
2. QC analysis Supplementary figure 27

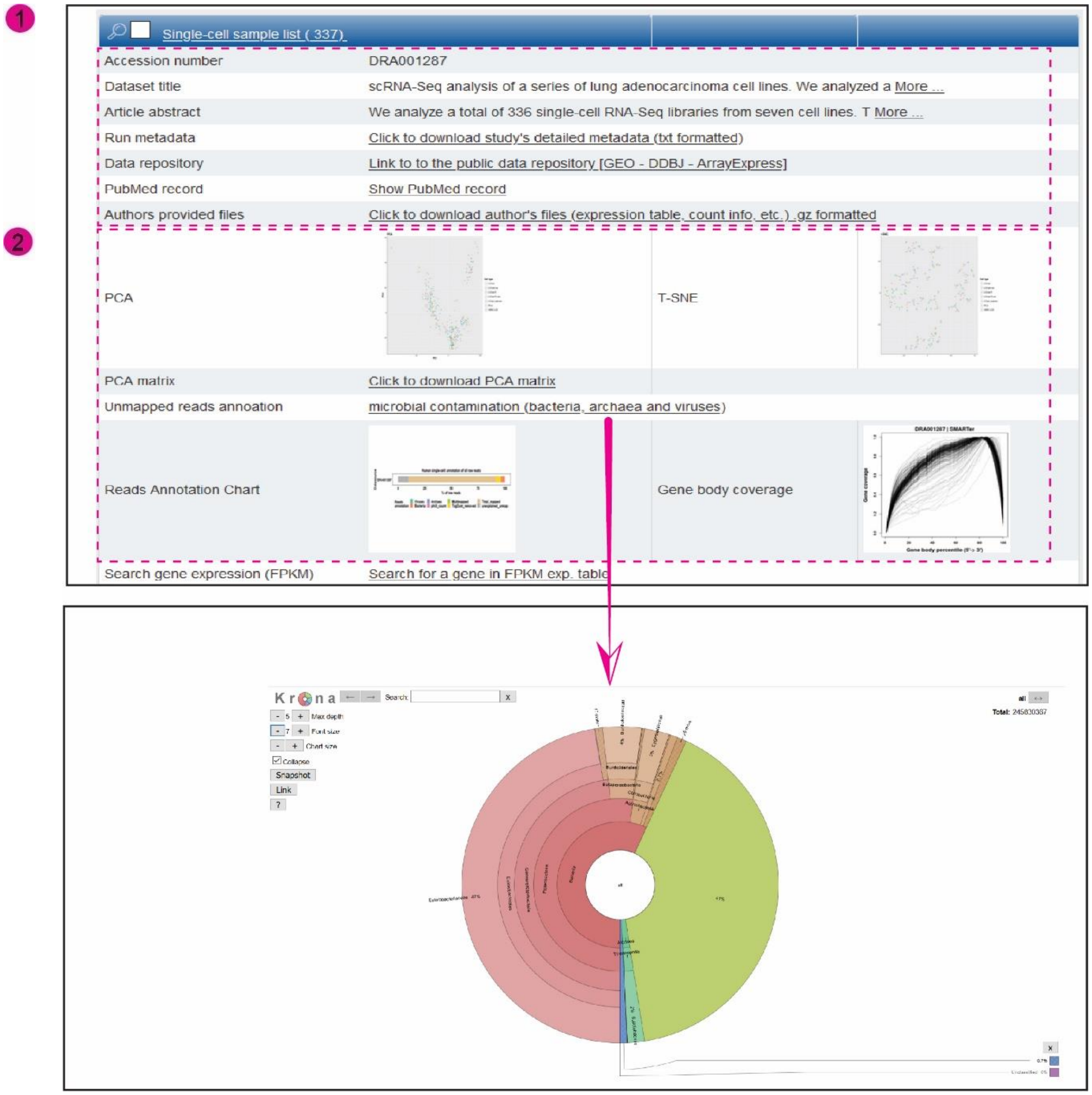


scRNA-seq dataset view. The top panel (1) lists basic attributes about the dataset and link to metadata files. The bottom panel of the screen (2) shows QC data and analysis results. The QC includes, PCA and t-SNE plots, Krona plot of the unmapped sequence reads annotation, bar chart of the sequence reads annotation and a gene body coverage plots.

3 | *Single-cell sample list*

This view provides detailed sample information, link to raw sequence files (FASTQ), BAM files, ration of intergenic expression, cell type annotation, library information etc.

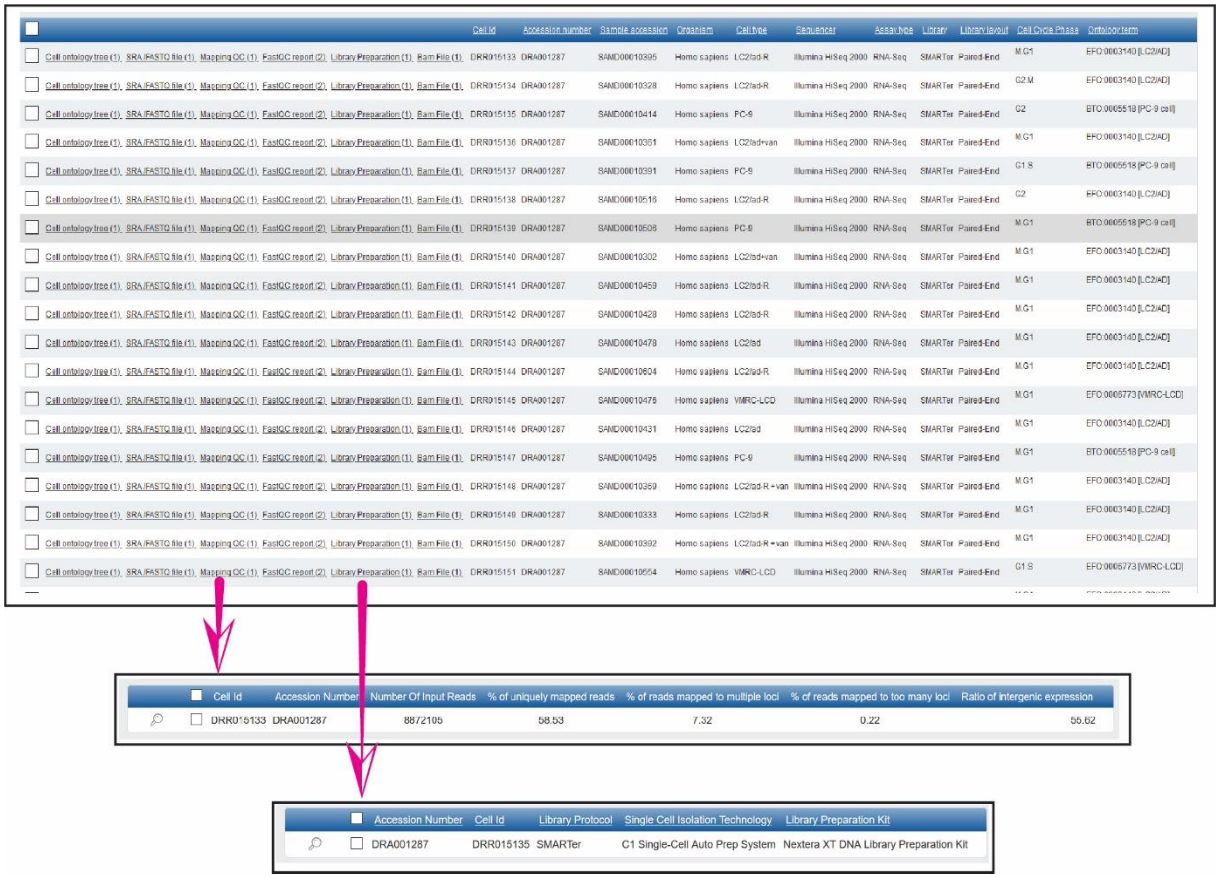


Single-cell sample, this view displays single-cell level metadata, cell annotation, cell-cycle phase, mapping QC (ratio of intergenic expression) and associated files (SRA file, BAM file.

4 | *Batch download*

SCPortalen provides batch download of the following files per each scRNA-seq dataset:

- BAM files
- FASTQC file
- Gene body coverage
- TPM and FPKM expression tables
- Unmapped reads analysis raw files

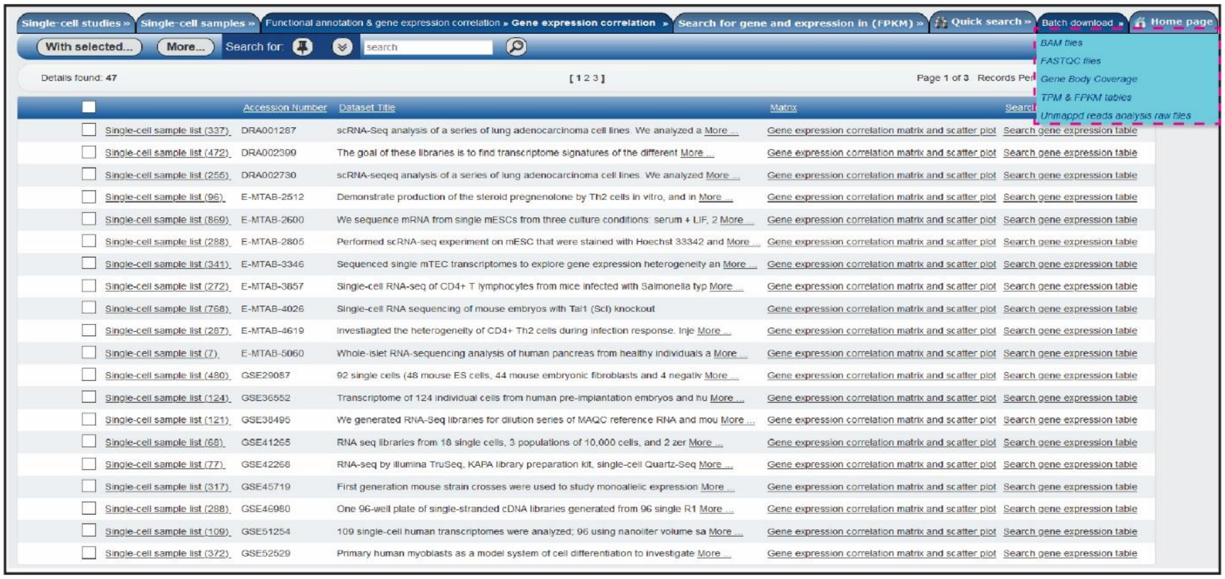

Batch download view of SCPortalen
